# On the Genes, Genealogies, and Geographies of Quebec

**DOI:** 10.1101/2022.07.20.500680

**Authors:** Luke Anderson-Trocmé, Dominic Nelson, Shadi Zabad, Alex Diaz-Papkovich, Nikolas Baya, Mathilde Touvier, Ben Jeffery, Christian Dina, Hélène Vézina, Jerome Kelleher, Simon Gravel

## Abstract

Population genetic models only provide coarse representations of real-world ancestry. We use a pedigree compiled from four million parish records and genotype data from 2,276 French and 20,451 French Canadian (FC) individuals, to finely model and trace FC ancestry through space and time. The loss of ancestral French population structure and the appearance of spatial and regional structure highlights a wide range of population expansion models. Geographic features shaped migrations throughout, and we find enrichments for migration, genetic and genealogical relatedness patterns within river networks across Quebec regions. Finally, we provide a freely accessible simulated whole-genome sequence dataset with spatiotemporal metadata for 1,426,749 individuals reflecting intricate FC population structure. Such realistic populations-scale simulations provide new opportunities to investigate population genetics at an unprecedented resolution.

**Lay Summary:** We all share common ancestors ranging from a couple generations ago to hundreds of thousands of years ago. The genetic differences between individuals today mostly depends on how closely related they are. The only problem is that the actual genealogies that relate all of us are often forgotten over time. Some geneticists have tried to come up with simple models of our shared ancestry but they don’t really explain the full, rich history of humanity. Our study uses a multi-institutional project in Quebec that has digitized parish records into a single unified genealogical database that dates back to the arrival of the first French settlers four hundred years ago. This genealogy traces the ancestry of millions of French-Canadian and we have used it to build a very high resolution genetic map. We used this genetic map to study in detail how certain historical events, and landscapes have influenced the genomes of French-Canadians today.

**One-Sentence Summary:** We present an accurate and high resolution spatiotemporal model of genetic variation in a founder population.

## Main Text

The tapestry of human genetic history is formed of ancestral lineages interwoven by generations of coalescence and recombination events (*1*). It was woven across geographic landscapes (*2*) by individual dispersal and historical waves of migrations that can sometimes be reconstructed by genomic analyses (*3, 4*). Yet the complex relationship between spatial migrations and genetic variation still poses formidable challenges (*5, 6*).

As a general trend, the limits of dispersal lead to continuous isolation-by-distance and a sometimes striking correlation between genetic and geographic distances (*7–9*). However, in any region, specific historical events or geographic barriers are often used to explain discrete patterns of population variation (e.g., (*5, 10, 11*)). Reconciling continuous variation into discrete “evolutionarily significant units” has proven to be difficult and sometimes misleading (*12–14*). While many studies have considered anisotropic migration models (*15*)), and even detailed models of geographic constraint or ‘resistance’ (*16, 17*), comparing these models to genetic data is challenging.

This study takes advantage of a population-scale spatially labelled pedigree (spatial pedigree) compiled from over four million Catholic parish records in the province of Quebec together with genotype data for 20,451 individuals, and new pedigree-aware simulation tools to provide a detailed spatiotemporal model of genetic variation at scales ranging from tens to thousands of kilometres. By including French and British individuals in our analyses we assess how much ancestral population structure has been preserved from these two populations. We highlight the relationship between river networks and genetic similarity as the past four centuries of European colonial history has been marked by rapid frontier expansion beginning along the shores of the St. Lawrence River, and eventually expanding up its tributaries. By tracing the ancestry of millions of individuals over space and time we describe a constellation of distinct founder events arranged along geographic features that defined transportation and economic activity. In doing so, we further bridge the gaps between family pedigrees and continental population structure as well as gaps between theoretical models and empirical demographic histories.

## Results

### Regional distribution of genetic variation

Quebec, a province in Canada, has a population of 8.6 million individuals, of which approximately 7.3 million speak French as a primary language. Because of strong historical correlation between Catholic religion, French ancestry, and French language, the genealogical records are much deeper and more complete for French Canadian (FC) individuals, by which we mean individuals who trace most of their ancestry to early French immigration irrespective of language. The majority of FC ancestry is derived from ∼ 8,500 settlers who migrated from France in the 17^th^ and 18^th^ centuries. The first 2,600 settlers contributed to two thirds of this gene pool (*18*). They occupied territory that had been inhabited and used by First Nations and Inuit peoples for thousands of years (*19*). Despite folk histories involving large amounts of indigenous ancestry among French Canadians (*20*), genetic and genealogical studies show that French Canadians born in Quebec carry on average less than 1% of ancestry tracing back to indigenous populations and the rest being mostly attributed to French ancestry (*21*).

Because pedigree data is much deeper and more complete among FC (*22*), we focused our genetic and genealogical analyses to 20,451 individuals inferred to be FC from the Cartagene (12,064 (*23*)) and Genizon cohorts (9,004, first reported here) (see Methods 3.1 for details on cohorts and ancestry inference). The genetic variation of this population was visualized using principal component analysis (PCA) (Fig. S1) and uniform manifold approximation and projection (UMAP) (Fig. 1A) (*24, 25*). A schematic summary of the entire FC spatial pedigree and the geographic location of 4,882 individuals linked to the spatial pedigree are shown in Fig. 1B and C. Visual inspection shows strong correlation of genetic and spatial proximity, and suggests that gradients in genetic variation coincide with geographical barriers and conduits like the St. Lawrence, Saguenay, and Chaudière Rivers or the Laurentian, and Appalachian Mountains.

**Figure 1:**
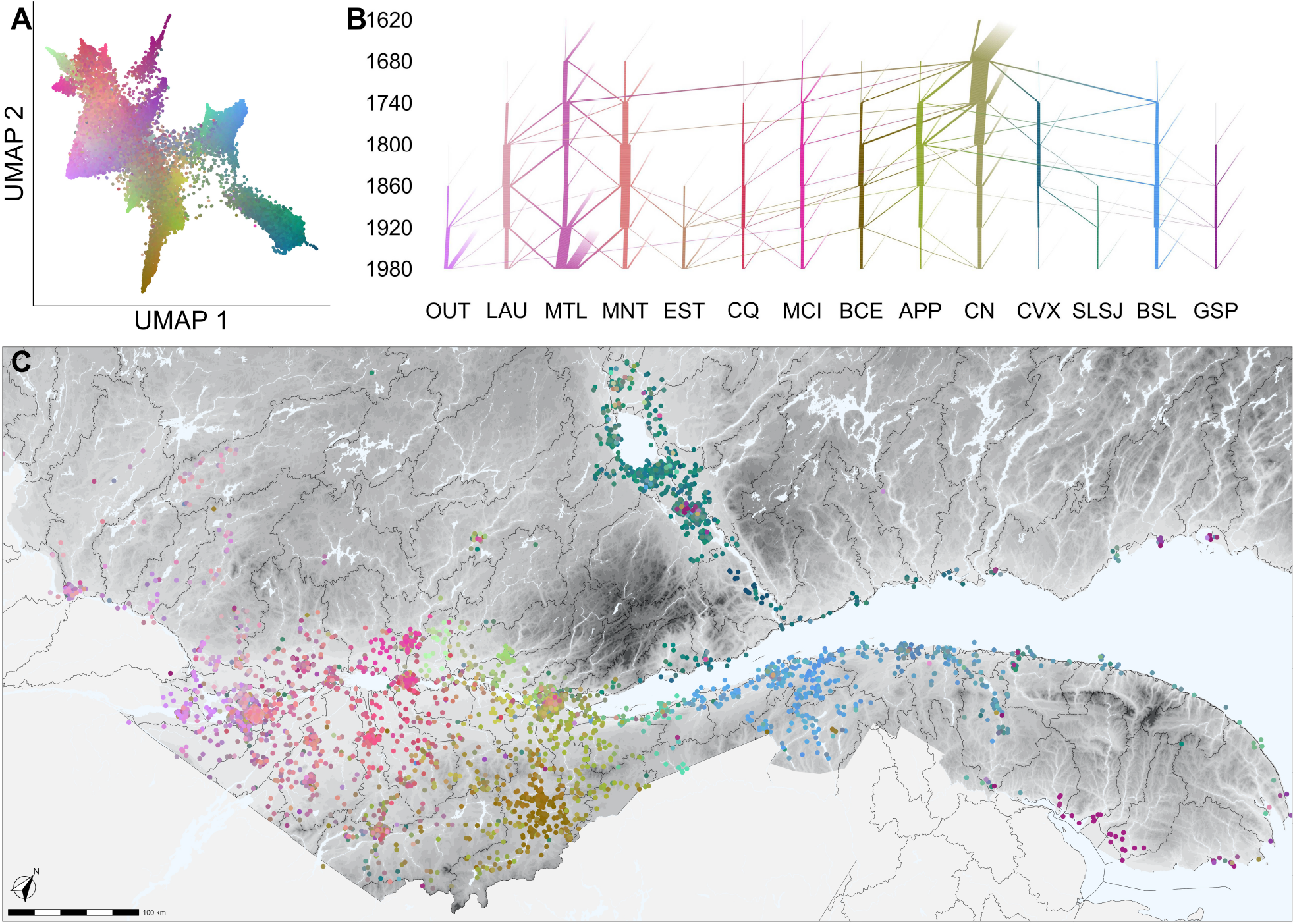
French Canadian genes and genealogies mirror Quebec’s geographies. (**A**) UMAP for 20,451 individuals with inferred French Canadian ancestry. Each individual is assigned a colour based on their location in a dimensionally reduced genetic projection space (see Supplementary Methods 3.2). (**B**) Visualizing the genetic ancestry of French Canadians across regions (x axis) over time of marriage (y axis). The thickness of the line a time *t* from location *A* to location *B* represents the amount of genetic material ancestral to the present day population. (see Supplementary Methods 4.4 for details). (**C**) Clusters in genetic space coincide with distinct geographic regions of Quebec defined by geographic features like rivers and mountains. Watershed boundaries are shown in black. Abbreviations: OUT – Outaouais; LAU – Laurentides; MTL – Montréal; MNT – Montérégie; EST – Estrie; CQ – Centre-du-Québec; MCI – Mauricie; BCE – Beauce; APP – Appalaches; CN – Capitale-Nationale; CVX – Charlevoix; SLSJ – Saguenay–Lac-Saint-Jean; BSL – Bas-Saint-Laurent; GSP – Gaspésie.

### French ancestry uprooted

Analyses of genealogical records show that a majority of French settlers came from regions in Northern and Northwestern France (Normandy, Ile-de-France, Aunis, Poitou, and Perche) (*26, 27*). However, successive waves of migration related to military and colonization objectives had different origins and demographics (*27*). To assess whether this history is reflected in the Quebec population structure, we compared the genomes of individuals living in different regions of Quebec and France. In agreement with historical records, FC share more recent ancestry with individuals from Northwestern France as measured by DNA that is identical by descent (IBD) (Fig. S2A). Furthermore, individuals living in the Outaouais and South Central regions of Quebec have lower rates of IBD with individuals from France (Fig. S2B). We used *F*_4_-statistics to assess how much French structure has been preserved and find no evidence of specific regions in Quebec being more similar to specific French and British counterparts (Fig. S3).This indicates that most present-day structure among FC is independent of any ancestral structure or to differential contributions by French and British founders.

### Simulated genomes with known transmission histories

To assess how much of the FC population structure can be accounted for by events following the arrival of French settlers, we generalized the msprime software (*28–30*) to perform genome-wide coalescent simulations conditioning on the known pedigree of the FC population. We defined the FixedPedigree ancestry simulation model in msprime version 1.2 to trace the ancestry of samples back through the genealogy accounting for coalescence and recombination (Fig. S4). To account for relatedness beyond the founders of the known pedigree, we modelled coalescence under a demographic model for European ancestry (*31*) (see Methods 2 for details).

We compared the simulations to ascertained data with a subset of 4,882 individuals who were both genotyped and linked to the pedigree. The PCA and UMAP of the simulations show clear qualitative agreement with the fine scale structure of ascertained genotype data (Fig. 2 and Fig. S3) as is evident from the first six PCs showing strong correlation (Fig. 2C). Thus, leading axes of genetic variation among FC reflect genetic drift that followed French settlement and is encoded in the spatial pedigree.

**Figure 2:**
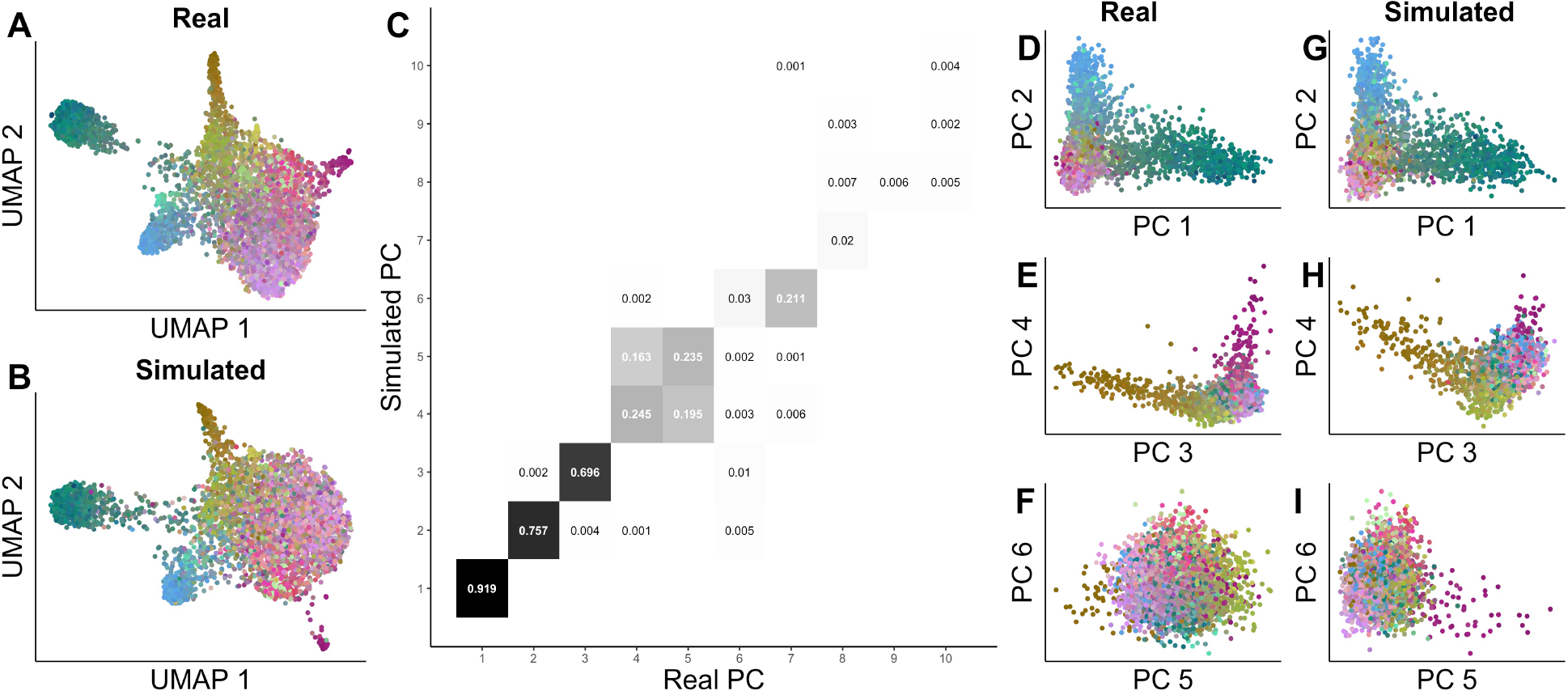
Simulated genomes capture observed population structure. Comparison of the same 4,882 individuals using simulated and observed genomes (coloured as in Fig. 1, see Methods 2 for details). (**A-B**) UMAP projections of observed and simulated genomes. (**C**) The correlation between observed and simulated principal components. (**D-F**) PCA projections of observed genomes. (**G-H**) PCA projections of simulated genomes.

Following this, we simulated whole genomes of 1.4M present day individuals with at least four grandparents linked to the pedigree. (see Fig. S5 for UMAP and PCA). Tree sequences of these simulations are freely available on Zenodo (https://doi.org/10.5281/zenodo.6839683). Although the tree sequences have been censored to remove personal identifying information, we have included temporal (decade) and spatial (latitude and longitude) information for the 1.4M samples and their ∼2M genealogically recoded genetic ancestors.

### Gene flow within watershed boundaries

Figure 1 shows axes of genetic variation that are restricted geographically, in a way more reminiscent of differentiation across alpine valleys (*5*) than of the continuous population structure seen in Europe (*9*). To assess how migration rates are impacted by topographical features like rivers and mountains, we visualize the major migration events shaping the FC population through space and time. Figure 3A exhibits waves of frontier expansion consisting of a series of sequential migrations up the tributaries of the St. Lawrence River ranging from tens to hundreds of kilometres in scale.

**Figure 3:**
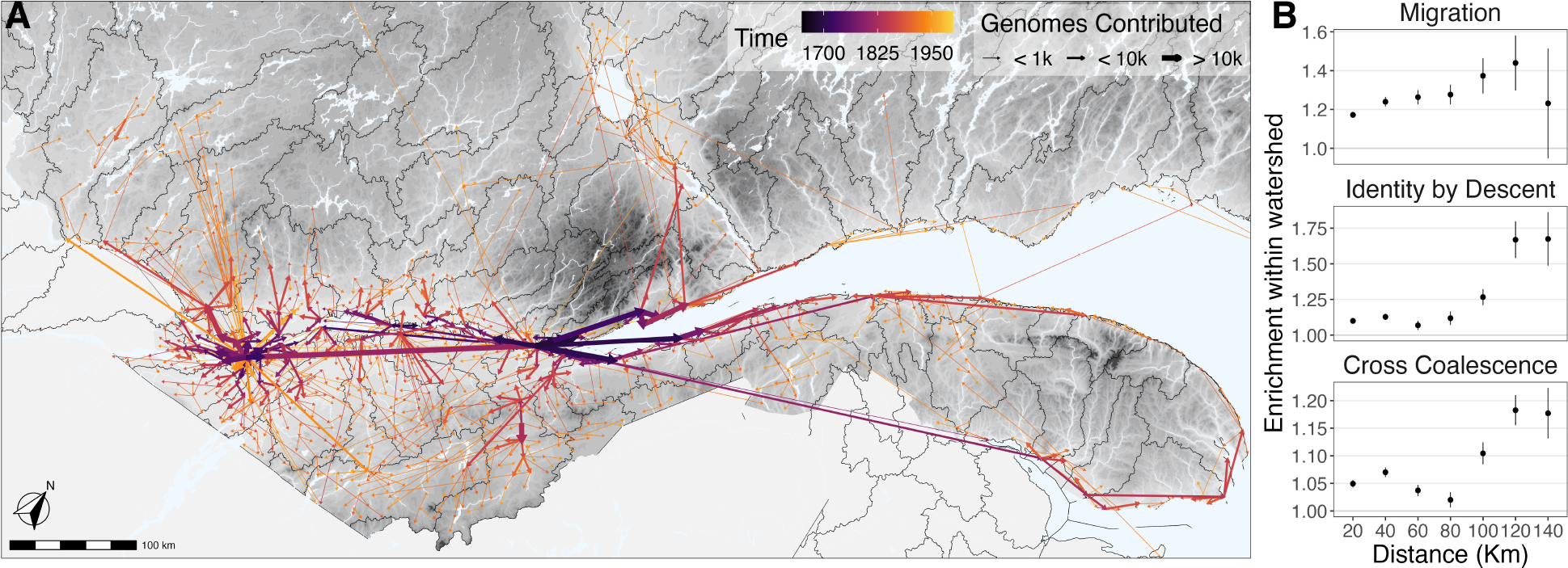
Watersheds influence French Canadian migrations and relatedness. (**A**) Primary routes of genetic ancestry. Segments link each town to the town from which migrants contributed the most genetic material. The width of segments indicates the total genetic contribution to present-day individuals and the colour indicates the mean date of when these contributions occurred historically. To avoid over plotting, we excluded the region of Abitibi-Témiscamingue and migrations of less than ten kilometres and migrations contributing less than ten genomes. (**B**) Relatedness is enriched within watershed. Black dots indicate the excess migrations and relatedness for towns within watershed relative to towns across watersheds at a fixed distance. See Methods 5 for details.

The first permanent French settlement took its name from the Algonquin word *kebec* making reference to the region where the St. Lawrence River becomes narrow (*32*). This was a strategic bridgehead location for the French as they sought to gain control of the main entrance to the Great Lakes in the 17^th^ century [(*33*), p 49–52]. Facing vast forested territory used and occupied by Iroquoian and Woodland First Nations (*19*), the French formed a fluvial colony with thin ribbons of settlements along shorelines using a riverfront land division strategy [(*33*), p 56–57].

To study patterns of relatedness and dispersal between individuals distributed in 1,698 parishes across the landscape, we considered three quantitative measures of relatedness and migration propensity between distinct geographic locations: migration rates, identity by descent, and cross-coalescence rates. All three show clear patterns of isolation by distance in all regions (Fig. S7). Given the importance of rivers in early settlement strategies, economic activity, and transportation (*34*), we hypothesized that relatedness and migration patterns broadly followed directions defined by local rivers and are therefore enriched for towns within watersheds. For each of the three metrics, we computed this enrichment for a tessellating set of 80 watersheds as a function of distance (Fig. 3B and Methods 5). We find an enrichment for the three metrics for all distances. At 120km, where the enrichment is strongest, we find a 40% enrichment in migration rates, 75% enrichment in genetic relatedness rates, and a 20% enrichment in cross coalescence rates. However, each region separately has a distinct migration pattern with varying degrees of topographical influence (Fig. 1C, Fig. S6).

### Historical migrations in space and time

To follow the formation of regional substructure in the FC population, we defined three regions with large proportions of individuals driving the top principal components (Fig. S8). We ascended the spatial pedigree for pairs of individuals living in each region, computed the realized kinship for each common ancestor (Methods 4.2.1), and determined the location, timing and stringency of historical genetic bottlenecks (Fig. 4 A-C).

**Figure 4:**
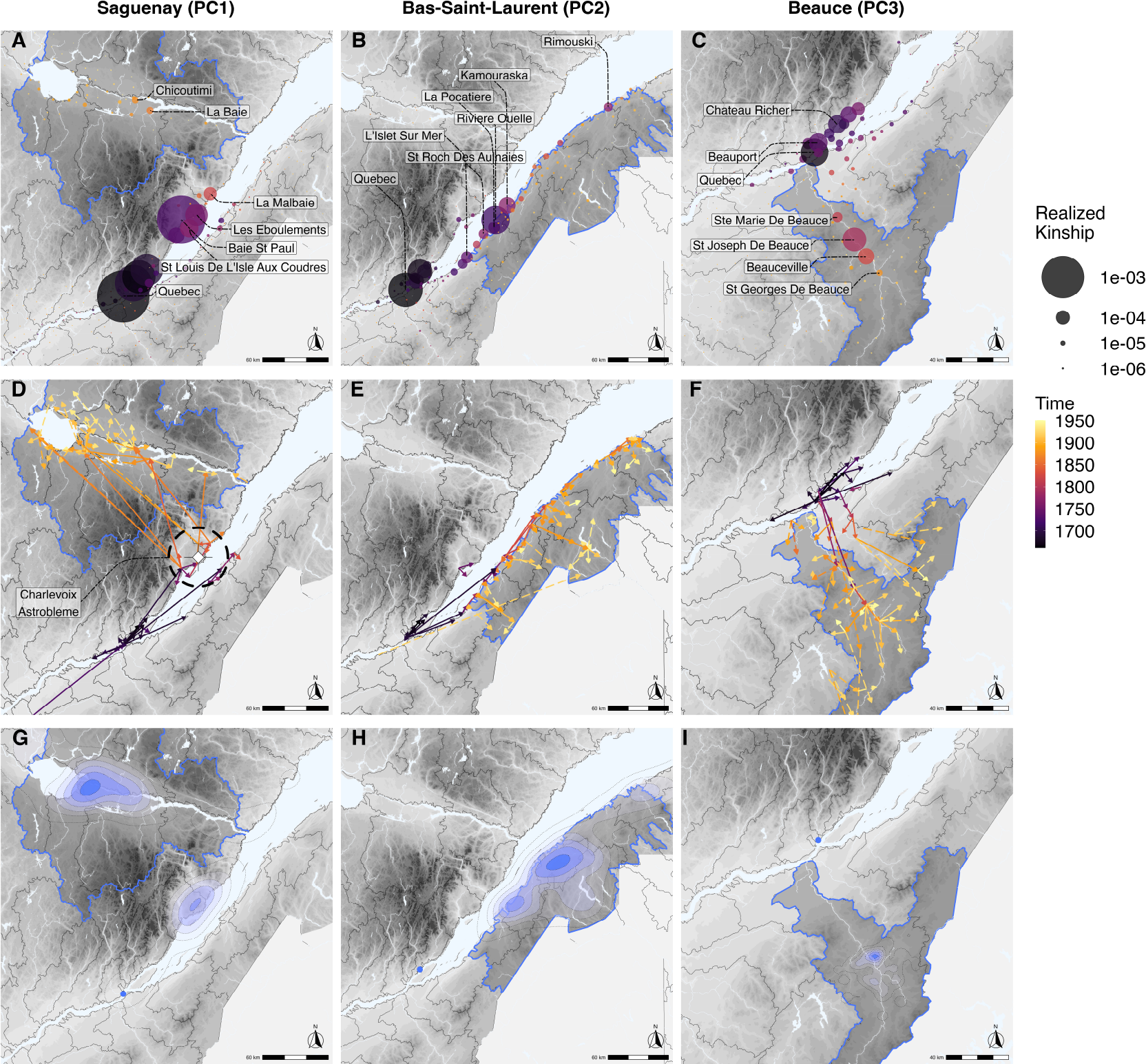
Historical migrations define population structure. (**A-C**) Time, location and stringency of genetic bottlenecks in three regions of Quebec defined by the top three principal components (Fig. S8). Size of points is proportional to realized kinship to present day FC population.Point color represents mean date of the realized kinship based on marriage dates of ancestors. (**D-F**) Major axes of migration as measured by the estimated genetic contributions to present day individuals living within highlighted regions. Dotted arrows indicate towns having the largest genetic contribution to towns within the highlighted regions, and solid arrows highlight the fifth percentile of migratory routes ranked by estimated genetic contribution. Arrow colours represent the mean time of the genetic contribution based on the marriage dates of ancestors. (**G-I**) The dispersal range for the single ancestor with the highest contribution to each of the three regions. For a given ancestor, we generate a heat map using the location of towns weighed by the average contribution of that ancestor to the probands of each town. The marriage location of each of the major contributors is indicated by a blue dot.

All regions exhibit substantial bottlenecks in and around Quebec city, but differ in their patterns of subsequent differentiation. The Saguenay-Lac-Saint-Jean (SLSJ) (Fig. 4 A,D) has a dominant early bottleneck in the region of Charlevoix, where an astrobleme (*35*) created a small pocket of fertile land within otherwise mountainous terrain (Fig. S10). Limited carrying capacity within the astrobleme led to demographic pressure and subsequent rapid expansion up the Saguenay River [(*33*), p 91], resulting in a vast majority of kinship predating the colonization of SLSJ. The Beauce region (Fig. 4 C,F) has a handful of bottlenecks in St-Joseph-De-Beauce and along the Chaudière River, with migrations reminiscent of a hub-and-spoke model. Finally, the Bas-Saint-Laurent (Fig. 4 B,E) has an assortment of bottlenecks including a dominant bottleneck in Rivière-Ouelle, but also more minor bottlenecks scattered across hundreds of kilometres of shoreline acting as a one dimensional regional hub for subsequent inland migrations (Fig. 4 E).

As expected in an expanding population, some early settlers (*super-founders*) had a large contribution to the population gene pool (*36, 37*). Here the top ten super-founders in each region contribute 32%, 11% and 12% of the realized kinship in SLSJ, Bas-Saint-Laurent and Beauce respectively (Fig. S11, S12, S13). And while each region has its deepest and most significant bottleneck near Quebec City, no two regions share the same super-founders (Fig. S14). To assess the overlap of each regional bottleneck, we computed cross-coalescence rates (*38*) (Methods 4.2.2 and Fig. S9) and find that 35% to 50% of kinship in Bas-Saint-Laurent and Beauce can be attributed to a shared bottleneck (Table 1), but the bottleneck in SLSJ is only 5% shared, reflecting much more intense founder events in Charlevoix and the region around Quebec City that are unique to SLSJ.

**Table 1:** Proportion of founder events shared between regions, as measured by the ration of cross-coalescence rate divided by within-region coalescence rate.

| Region | Proportion of bottleneck shared with: |  |  |
| --- | --- | --- | --- |
|  | SLSJ | Bas-Saint-Laurent | Beauce |
| SLSJ | - | 0.055 | 0.035 |
| Bas-Saint-Laurent | 0.456 | - | 0.358 |
| Beauce | 0.412 | 0.516 | - |

Not all regions of Quebec exhibit such spatially defined bottlenecks. Even though Abitibi-Témiscamingue (AT) was settled by a similar process of rapid frontier expansion as seen in SLSJ, it did not lead to bottlenecks reflected in leading axes of genetic differentiation. In contrast to the events in SLSJ, the settlers to AT came from numerous villages scattered throughout the province. While many of the villages in AT have measurable bottlenecks, at a regional level these bottlenecks seldom overlap. To illustrate this, we consider the villages of Remigny and Rollet separated by twenty kilometres along the Ottawa River (Fig. S15). Cross-coalescence rates of these villages have 11% overlap. For comparison, La Baie and Roberval in SLSJ have 70% overlap despite being over one hundred kilometres apart. Even though towns in AT show limited evidence of a shared founder effect, the parallel founding events create sub-structure beyond isolation-by-distance (*39*).

## Discussion

Classical population genetic models often approximate reproduction and mate selection as a uniform random process. By approximating the effects of a myriad of individual motivations and choices that are unavailable to scientists, classical models provide an explanation for trends such as drift and selection. However, large population samples highlight the limitations of these models (*4, 40*). As we give up the simplified assumption of uniform random mating, the number of demographic parameters relevant to evolution grows rapidly.

The BALSAC pedigree – a particularly complete population scale spatial pedigree – has been instrumental in identifying multigenerational demographic effects like reproductive advantages of being on a wave-front expansion (*36*) or the transmissibility of family size (*41*) and migration propensity (*42*). Others have used it to study variation in runs of homozygosity (*43*) and its historical determinants, such as the kinship of the first settlers of Charlevoix (*34*) or the delay in the settlement of the Saguenay due to restrictions related to the fur trade [(*33*), p 91].

Here we sought to develop a comprehensive genetic model that captured all these effects, and more. We used this model to highlight how one of the best-studied human founder populations in SLSJ was influenced by the unique geography of the region that was shaped by a cosmic event occurring 400Mya in Charlevoix. We also found concordant genetic and genealogical support for the idea that geographic features like rivers and mountains played a systematic role in influencing migration rates and defining major axes of genetic variation. Finally, the strong correlation between empirical and simulated genetic data provides evidence that the structure within the FC population can largely be attributed to events in North America, while the population kept a genetic signature of the regions in France contributing more early French settlers.

We describe how geological, social, historical effects, as well as idiosyncratic events, translated in genetic variation patterns at various geographic scales and over centuries. Although our simulations are based on a real pedigree, they do not contain identifying information and we can freely share this genome-wide dataset along with spatiotemporal metadata for over 1.4M individuals. Of course, these simulations are far from perfect. They do not account for natural selection beyond what is captured in the pedigree (*36*). The pedigree itself contains some amount of recording errors. Mismatches between genealogical ancestry and biological ancestry are not uncommon, and these will be difficult to overcome (*22, 40*). However, we believe that the genetic model, the simulation tools, and the publicly available simulated data we described provide a lens to investigate population genetics at an unprecedented resolution.

## Acknowledgements

We acknowledge the contributions of Ivan Krukov who we consider eligible for authorship, but were unable to contact for approval. We are grateful to all of the participants who enabled this study by contributing their DNA, and to the participants who provided family information enabling the reconstruction of their genealogy. For data from Quebec, we thank the BALSAC team for their management and curation of the genealogy database, the CARTaGENE team and the Genome Quebec for their management and curation of genotype data. For data from France, we thank the EREN team for their management and curation of SU.VI.MAX genotype data. We thank Wilder Wohns, Yan Wong, and Gil McVean for enlightening conversations at early stages of the project, and Aaron Ragsdale for his comments on early drafts of this manuscript.

## Author contributions

Conceptualization: LAT, DN, CD, HV, JK, SG

Data curation: LAT, BALSAC, CARTaGENE, Genome Quebec, EREN, MT

Formal analysis: LAT

Funding acquisition: LAT, JK, SG

Investigation: LAT, SG

Methodology: LAT, DN, SZ, ADP, JK, SG

Project administration: LAT, JK, SG

Resources: SG

Software: LAT, DN, IK, SZ, NB, BJ, JK, SG

Supervision: JK, HV, SG

Validation: LAT, DN, IK, NB, BJ, JK, SG

Visualization: LAT

Writing – original draft: LAT

Writing – review & editing: LAT, AR, CD, ADP, SZ, JK, HV, SG

## Data and materials availability

R code for analyses and visualizations are available (https://github.com/LukeAndersonTrocme/genes_in_space/). Tree sequences of simulated genomes are freely available on Zenodo (https://doi.org/10.5281/zenodo.6839683). Quebec genotype data are available upon request to (www.cartagene.qc.ca) and (https://www.mcgillgenomecentre.ca/) for CARTaGENE and Genizon cohorts respectively. Genealogical data are available on the Scholars Portal Dataverse platform from the University of Québec in Chicoutimi (https://doi.org/10.5683/SP3/BW7DIG.)

