## Supporting Materials for "On the Genes, Genealogies, and Geographies of Quebec"

**5 Enrichment analysis** **32**

**List of Symbols** **36**

**Supporting tables and figures** **37**

### 1 Data

#### 1.1 Genetic data

The genotype data used in this study was compiled from three separate cohorts, each of which was imputed separately using the Michigan Imputation Server then merged (Supplementary Figure S17). The regional distribution of the samples included in the study is summarized in Figure 1. The Genizon cohort is comprised of 9,961 genotyped individuals from Quebec of which 2,431 of them consented to and were successfully linked to genealogical records. The genotype data from this cohort was produced on 4 different chips (HumanHap375, HumanHap550, Illumina1M and Human610-Quad) and due to data being unavailable for chromosome 22 for one of the Genizon chips, we restricted all of our genetic analyses to the first twenty one chromosomes. The CARTaGENE dataset (`www.cartagene.qc.ca`) used here comprises of 12,062 genotyped individuals from Quebec of which 5,733 consented to and were successfully linked to genealogical records. The genotype data from this cohort was also produced on 4 different chips (Omni2.5, Axiom2.0, GSAv1 and GSAv2). The SUVIMAX cohort includes 2,184 genotyped individuals from France, see (*11*) for details. For details about the downsampled sequence data from the 1000 Genomes GBR population, see (*44*).

#### 1.2 Genealogical data

BALSAC is a comprehensive genealogy of the French-Canadian population of Quebec compiled from 4,282,960 marriage records dating back to the $17^{\text{th}}$ Century (*22*). Data are available on the Scholars Portal Dataverse platform from the University of Québec in Chicoutimi (*45*).

#### 1.3 Geographical data

Layers of geographical data including rivers, lakes, watersheds, provincial, and federal boundaries were downloaded from the government of Canada geobase `https://open.canada.ca/`.

Polygons from each layer were simplified using ms_simplify function from the rmapshaper R library (*46*) and projected onto the EPSG:4326 coordinate system. Digital elevation model GeoTIF files were downloaded from the government of Canada `https://open.canada.ca/`. The hydrological and altimetry data are licensed under the Open Government Licence allowing for their use, modification and publication.

##### 1.4 Demographic data

Population size estimates of the province of Quebec were obtained from the preliminary results of the 2020 Census (*47*) and estimates of the number of individuals who speak French as a primary language from (*48*).

#### 2 Genome simulations

##### 2.1 Fixed pedigree simulation model

We extended the `msprime` software to include support for simulations conditional on a fixed pedigree. This new extension simulates the effects of recombination and the transfer of ancestral material from children to parents based on the structure of the pedigree. This simulation model can use user specified recombination rates (or maps) and accounts for founders living at different times and accommodates founders from multiple source populations. Ancestry beyond the fixed pedigree can be simulated using arbitrarily complex demographic models including those specified in the PopSim Consortium (`https://popsim-consortium.github.io/stdpopsim-docs/stable/index.html`) (*49*).

##### 2.2 Model specifications

Even though our software implementation accommodates multiple source populations for the founders, we considered a single source population of European ancestry as defined by the two population out-of-Africa model of Tennessen et al. (*31*). The chromosome length and recombination rate for each simulated chromosome was defined by the GRCh37 hapmapII genetic map (*50*). The genomic regions belonging to centromeres and telomeres were excluded from our simulations given their lack of documented recombination events. Once the tree sequences were constructed, mutations were added to branches of the tree at a rate of $3.62 \times 10^{-8}$ per basepair per meiosis. This unusually high mutation rate is chosen to match the mutation rate $\mu_{cds} = 2.35 \times 10^{-8}$ used in (*31*) to infer the demographic model, while accounting for the difference in diversity between coding and non-coding sequences. Because the mutation rate varies along the genome, we scaled this mutation rate to match genome-wide expectations using the relative rate of intergenic to coding polymorphism, i.e., $\mu_{int}/\mu_{cds} = 1.53$ as described in (*51*). A summary of model specifications are in table S3.

A GitHub repository with code to run the genome simulation pipeline is available (`https://github.com/LukeAndersonTrocme/genome_simulations`) and extensive documentation of Msprime (`https://tskit.dev/msprime/docs/latest/api.html#msprime.FixedPedigree`).

#### 2.3 Comparing simulations to ascertained data

To compare the simulated genomes to ascertained genotype data, we downsampled the simulations to match the density of the genotype data. We did so by removing variants below a 5% minor allele frequency and linkage disequilibrium pruning such that both datasets contained $\sim$60,000 variants. We restricted our simulations to 4,882 individuals from a total of 5,402 individuals who consented to be linked to the FC pedigree. We excluded 100 individuals who did not have all four grandparents present in the pedigree. We also excluded 420 individuals with second cousins or closer relatives to match a quality control step typical in population genetic studies. We performed a principal component analysis (*52*) on both ascertained and simulated data using the same 4,882 individuals, and for visualization purposes, we used the same three dimensional colours used in Figure 1.

### 3 Genetic statistics

#### 3.1 Genetic ancestry of French Canadians

The majority of individuals in Quebec derive FC ancestry (*47, 48*), as such we expect the majority of the participants in our cohorts also derive FC ancestry despite only a fraction of them being linked to the genealogy. For the purposes of visualizing the population structure of FC in Figure 1A and C, we sought to leverage the large number of FC participants included in our PCA and UMAP analyses. We defined a threshold based on the genotype data from participants linked to this genealogy and their projections along the first principal component. This threshold kept 21,146 genotyped individuals with presumed FC ancestry, and excluded 617 individuals from our analyses (see Supplementary Figure S18 and Supplementary Table S1).

#### 3.2 Dimension reduction

Flashpca2 was used to performed our principal component analysis (PCA) (*52*) to generate (Supplementary Figure S1) and the R package uwot (*53*) for our uniform manifold and approximation projection (UMAP) (*24*) used to generate Figure 1A. This method takes the first ten principal components of genetic data as input and reduces this high dimensional data to a lower dimension while seeking to preserve local neighbourhoods.

The colours used in Figure 1 were determined by reducing the top ten principal components of genotype data to a three dimensional UMAP and then converted each $x, y, z$ coordinate into an RGB value that is unique to each individual (*54*).

#### 3.3 F statistics

F statistics used in Supplementary Figure S2A were computed using the admixture R package (*55, 56*). We used all 2,184 genotyped individuals from France as they all had regional geospatial information available. We also included 94 British individuals from the 1000 Genomes Project as an additional potential founding population. Together, we refer to the British and French samples as European. For the French Canadian samples, we used the 4,882 individuals linked to the genealogy with geospatial information. The population groupings used for the French Canadian individuals were watershed boundaries, the French groupings used seven French regions (South-East, South-West, West, North-West, Central, Isle-of-France, East) (Table S2), and the British GBR individuals were kept in a single group.

We computed all pairwise $F_4$ statistics $F_4(qc1, qc2, eu1, eu2)$ between French and Quebec regions. As a positive control we also computed the complement $F_4(qc1, eu1, qc2, eu2)$ statistics for all regions.

#### 3.4 Identity by descent

Using the genotype data from samples from Quebec and from France, in the Genizon, CARTaGENE, SUVIMAX, and GBR cohorts, we first phased the data using Shapeit4 (*57*) and downsampled to a set of common SNPs before computing pairwise IBD using Hap-IBD for all samples (*58*) using a minimum segment length of seven centimorgans.

We used all 2,184 genotyped individuals from France as they all had regional geospatial information available. For the French Canadian samples, we used the 4,882 individuals linked to the genealogy with geospatial information. The population groupings used for the French Canadian samples were geographic regions as defined in Figure 4 and the French groupings used the seven French regions (South-East, South-West, West, North-West, Central, Isle-of-France, East).

Using the rates of IBD between all pairs of individuals, we computed the average IBD sharing rates $g(A, B)$ between sets $P^A$ and $P^B$ of individuals in towns $A$ and $B$ respectively:

$$g(A, B) = \frac{1}{|P^A||P^B|} \sum_{i \in P^A, j \in P^B} IBD(i, j) \quad \text{(S1)}$$

where $IBD(i, j)$ is the total length in centiMorgans of IBD segments between individuals $i$ and $j$, $|P^A|$ and $|P^B|$ are the sample sizes in towns $A$ and $B$.

### 4 Genealogical statistics

#### 4.1 Estimated contributions

Let us call $\mathrm{K}^P(i)$, the expected genetic contributions (i.e., the length of inherited genetic material in centiMorgans) of an individual $i$ to a set $P$ of probands,

$$\mathrm{K}^P(i) = \sum_{p \in P} \mathrm{K}^p(i) \tag{S2}$$

where $\mathrm{K}^p(i)$ is the expected contribution of individual $i$ to proband $p$. Similarly, the contributions $\mathrm{K}^P(I)$ of a set $I$ of individuals to probands $P$ is simply

$$\mathrm{K}^P(I) = \sum_{i \in I} \mathrm{K}^P(i) \tag{S3}$$

#### 4.2 Coalescence rates

##### 4.2.1 Within-population coalescence

Founder effects and genetic bottlenecks result in excess kinship among individuals from a population. Given a spatial pedigree, we can identify the specific common ancestors that contribute to kinship between any two individuals, and therefore track a founder effect in space and time. Given a set of probands $P$, define $\lambda^P(i)$ as the total expected pairwise kinship realized in individual $i$. In other words, $\lambda^P(i)$ measures how often $i$ is the most recent common ancestors of pairs of individuals in $P$. This can be estimated rapidly from the genetic contributions $\mathrm{K}^P(\cdot)$ of the different offspring to individual $i$:

$$\lambda^P(i) \approx \frac{1}{4} \sum_{(m,n)} \mathrm{K}^P(m)\mathrm{K}^P(n) \tag{S4}$$

where $(m, n)$ are the ordered pairs of offspring for individual $i$.

##### 4.2.2 Cross coalescence

In a similar fashion as the within-region realized kinship described above, we can compute the cross coalescence rate between regions to identify the specific common ancestors that contribute to kinship between any two individuals in *different* regions, and therefore determine whether certain founder effects are shared between regions.

Given a set of probands $P^A$ and $P^B$, define $\lambda^{P^A P^B}(i)$ as the total expected pairwise kinship realized in individual $i$. In other words, $\lambda^{P^A P^B}(i)$ measures how often $i$ is the most recent common ancestors of pairs of individuals in $P^A$ and $P^B$. This can be estimated rapidly from the genetic contributions $\mathrm{K}^{P^A}(\cdot)$ and $\mathrm{K}^{P^B}(\cdot)$ of offspring to individual $i$:

$$\lambda^{P^A P^B}(i) \approx \frac{1}{4} \sum_{(m,n)} \mathrm{K}^{P^A}(m)\mathrm{K}^{P^B}(n) \tag{S5}$$

where $(m, n)$ are the ordered pairs of offspring for individual $i$.

##### 4.2.3 Overlap of relative cross coalescence

To assess the percent overlap of two bottlenecks, we can contrast the cross-coalescence rates to the within-population coalescence rates. Given a set of probands $P^A$ and $P^B$, we define $\gamma^{P^A P^B}$ as the ratio of cross coalescence to within-population coalescence $P^A$ and $P^B$ (*38*):

$$\gamma^{P^A P^B} = \frac{2\Lambda^{P^A P^B}}{\Lambda^{P^A} + \Lambda^{P^B}} \tag{S6}$$

where

$$\Lambda^{P^A P^B} = \frac{1}{|P^A||P^B|} \sum_i \lambda^{P^A P^B}(i),$$

$$\Lambda^{P^A} = \frac{2}{|P^A| \times (|P^A| - 1)} \sum_i \lambda^{P^A}(i),$$

and

$$\Lambda^{P^B} = \frac{2}{|P^B| \times (|P^B| - 1)} \sum_i \lambda^{P^B}(i).$$

###### 4.2.4 Normalization

The relative cross-coalescence rate can be interpreted as a measure of the similarity of coalescence history between pairs of individuals within and across population, and is therefore normalized by sample size. This is shown, for example, in Figure 3. By contrast, when trying to compare the amount of kinship realized in historical individuals, we want to account for the fact that individuals who had many desendents contributed a lot to present-day kinship. In Figure 4, we therefore use a non-normalized kinship measure to identify the total contributions of individuals to present-day relatedness.

##### 4.3 Migration rates

For the purposes of studying historical migrations, we assume that individuals are born in the location where their parents married, and migrate to the location of their own marriage. We note that this definition is incomplete as it does not account for a cultural practice within French-Canadians where couples would tend to marry in the local church of the female counterpart and then move to the region of origin of the male counterpart. In our analyses, our estimates of migration rates are averaged across both sexes and all generations.

###### 4.3.1 Contribution date

We defined above the genetic contribution for a set of individuals. To study the contributions of migrants specifically, we consider the set $M_{a \to b}$ of individuals born in source-town $a$ and married in sink-town $b$ and their total contribution $\mathrm{K}^P(M_{a \to b})$. To characterize the time period where contributing migrations occurred, we also report the mean contribution date $d(M_{a \to b})$ defined as

$$d(M_{a \to b}) = \sum_{i \in I_{a \to b}} w_i d_i \quad \text{(S7)}$$

where $d_i$ is the marriage date of individual $i$ and $w_i$ is a weight proportional to the total estimated genetic contribution $\mathrm{K}^P(i)$.

##### 4.3.2 Relative emigration rate

In population genetics, a commonly used definition of migration rate is the fraction of immigrants over the total population size. However, in our enrichment analysis, we seek to compare multiple *inbound* migration rates $\delta_{a\to b}$ from multiple choices for source-town $a$ to the same reference sink-town $b$. For this reason, we use a less common definition of migration rate of :

$$\delta_{a\to b} = \frac{|M_{a\to b}|}{N_a} \tag{S8}$$

where $N_a$ is the number of people born in source-town $a$.

The *emigration* rates account for the different population sizes of different source-towns since we normalize over $N_a$ rather than $N_b$. The sum of migration rates $\delta_{A\to b}$ for a set $A$ of source-towns to $b$.

$$\delta_{A\to b} = \sum_{a\in A} \delta_{a\to b}. \tag{S9}$$

#### 4.4 Genealogy flow plot

We generated a visual summary of French Canadian ancestry using the riverplot R package. The x axis of the plot was generated by grouping together individuals based on administrative region boundaries in Quebec with some slight modifications highlighted in Supplementary Figure S16. The line thickness was obtained by aggregating the total estimated genetic contributions $\mathrm{K}^P(I_{a\to b,t})$ of individuals $I_{a\to b,t}$ to all probands $P$ in the genealogy based on where they were born $a$, where they married $b$, and when they married $t$. This plot includes missing data as contributions fading to white for each region and time bin. To avoid overplotting, we exclude small migrations contributing less than the top 20 percent of migrant contributions. The y axis of the plot is separated into 60 year time bins starting from 1620 and ending in 1980. A small number of individuals in this dataset married either before or after this time range were added to the first and last time bins respectively.

##### 4.5 Code availability

The R code used to compute cross coalescence and visualize the genealogy are available here `https://github.com/LukeAndersonTrocme/genes_in_space/tree/main/supplementary` `code`.

### 5 Enrichment analysis

Using three metrics measuring the relatedness of individuals in a pair of towns – migrations, identity by descent, and cross coalescence – this enrichment analysis tests the null hypothesis that pairs of towns at a given distance have equivalent relatedness rates regardless of whether they share a watershed. To compare sets of towns of equal distance, we define $T$ to be a set of distal towns within a twenty kilometre wide annulus whose inner radius is $d$ kilometres from a reference town $b$ and $S$ be a set of distal towns within the same watershed as $b$. The distal towns that are in the intersection of the sets T and S (i.e. they are within the same watershed as $b$ and within the annulus defined by $d$)

$$T' = T \cap S. \tag{S10}$$

From this, we can define the baseline of our enrichment as the fraction of distal towns – with sampled individuals – sharing a watershed with reference town $b$ within a fixed distance $d$

$$c(b) = \frac{|T'|}{|T|}, \tag{S11}$$

For each of the three statistics considered for watershed enrichment, we define

$$\omega_m(b, a) \tag{S12}$$

as the value of metric $m$ between reference towns $b$ and distal towns $a$. The sum of $\omega_m(b, a)$ over sets of distal towns $T'$ and $T$ are

$$\Omega_m(b, T') = \sum_{a \in T'} \omega_m(b, a), \tag{S13}$$

and

$$\Omega_m(b, T) = \sum_{a \in T} \omega_m(b, a), \tag{S14}$$

respectively.

From this, we define the fraction $\eta_m(b)$ of $\Omega_m(b)$ distal towns sharing a watershed with reference town $b$ within a fixed distance $d$ as

$$\eta_m(b) = \frac{\Omega_m(b, T')}{\Omega_m(b, T)}. \tag{S15}$$

Finally, we define the enrichment of metric $m$ for distal towns sharing a watershed with reference town $b$ within a fixed distance $d$ as

$$\epsilon_m(b) = \frac{\eta_m(b)}{c(b)}. \tag{S16}$$

**Example**

To illustrate our enrichment metric, let us consider a null model where

$$\omega_m(b, t) = 1$$

for all pairs of distal towns $t$ and reference towns $b$. In this case,

$$\Omega_m(b, T') = T'$$

and

$$\Omega_m(b, T) = T,$$

where

$$\eta_m(b) = \frac{T'}{T} = c(b),$$

which yields

$$\epsilon(b) = \frac{\eta_m(b)}{c(b)} = 1.$$

**Enrichment metrics**

As mentioned in the section above, the three metrics used in our enrichment analysis are IBD, migrations, and cross coalescence.

The average length of DNA that is IBD between individuals in a given town $a$ and a reference town $b$ similar to 3.4:

$$\omega_{IBD}(b,a) \equiv g(a,b) = \frac{1}{|P^a||P^b|} \sum_{i \in P^a, j \in P^b} IBD(i,j) \tag{S17}$$

where $IBD(i,j)$ is the total length in centiMorgans of IBD segments between individuals $i$ and $j$ in towns $a$ and $b$ respectively.

The emigration rate from a given town $a$ to a reference town $b$ defined in 4.3.2 :

$$\omega_{mig}(b,a) \equiv \delta_{a \to b} = \frac{|M_{a \to b}|}{N_a} \tag{S18}$$

where $N_a$ are the number of people born in distal town $a$.

The relative cross coalescence between individuals in a given distal town $a$ and a reference town $b$ defined in 4.2.3 :

$$\omega_{coal}(b,a) \equiv \gamma^{ba} = \frac{2\Lambda^{ba}}{\Lambda^b + \Lambda^a} \tag{S19}$$

where $\Lambda^{ba}$ is the ratio of relative cross coalescence and $\Lambda^b$ and $\Lambda^a$ are the relative within-population coalescences for probands in $b$ and $a$ respectively.

**Implementation**

Because the Canadian National Hydrographic Network classifies the hundreds of kilometres of shoreline of the St. Lawrence River as one single watershed, we excluded this unusual watershed from the analysis by removing all reference towns within this watershed. We note that distal towns within this watershed remain in the analysis.

### List of Symbols

| Symbol | Definition | Obs. Exp. | Unit |
|---|---|---|---|
| $IBD(i,j)$ | length of IBD segments between individuals $i$ and $j$ | O | centiMorgans |
| $g(A,B)$ | mean IBD sharing between pairs of individuals in towns $A$ and $B$ | O | centiMorgans |
| $P^A$ | probands in town $A$ | O | set |
| $\lvert P^A\rvert$ | sample size in town $A$ | O | - |
| $\mathrm{K}^p(i)$ | genetic contribution of individual $i$ to proband $p$ | E | genomes |
| $\mathrm{K}^P(i)$ | total genetic contributions of individual $i$ to proband set $P$ | E | genomes |
| $\mathrm{K}^P(I)$ | total contributions of a set $I$ of individuals to proband set $P$ | E | genomes |
| $\lambda^P(i)$ | kinship realized in individual $i$ given a set of probands $P$ | E | probability for time of first coalescence for all pairs of lineages |
| $\lambda^{AB}(i)$ | kinship realized in individual $i$ given sets of probands $A$ and $B$ | E | probability for time of first coalescence for all pairs of lineages |
| $\gamma^{AB}$ | ratio of cross-coalescence to within-population coalescence of sets of probands $P^A$ and $P^B$ | E | probability for time of first coalescence for all pairs of lineages |
| $\Lambda^{AB}$ | cross-coalescence rate of the sets of probands $P^A$ and $P^B$ | E | probability for time of first coalescence for all pairs of lineages |
| $\Lambda^A$ | within-population coalescence rate of set of probands $P^A$ | E | probability for time of first coalescence for all pairs of lineages |
| $M_{a\to b}$ | set of individuals born in source-town $a$ and married in sink-town $b$ | O | set |
| $\mathrm{K}^P(M_{a\to b})$ | genetic contribution of individuals born in $a$ and married in $b$ | E | genomes |
| $d(M_{a\to b})$ | mean contribution year of individuals born in $a$ and married in $b$ | E | years |
| $\lvert M_{a\to b}\rvert$ | number of migrants from $a$ to $b$ | O | individuals |
| $N_a$ | number of individuals born in $a$ | O | individuals |
| $\delta_{A\to b}$ | inbound migration rate of set $A$ of source-towns to sink-town $b$ | O | proportion |
| $T$ | towns within a defined annulus centred on town $b$ | O | set |
| $S$ | towns within the same watershed as $b$ | O | set |
| $T'$ | towns within the same watershed and annulus centred on town $b$ : $T\cap S$ | O | set |
| $c(b)$ | fraction of towns sharing a watershed with $b$ within a defined annulus : $\frac{\lvert T'\rvert}{\lvert T\rvert}$ | O | proportion |
| $m$ | metric whose enrichment is being considered (i.e. migration rate, IBD sharing rate, cross coalescence rate) | - | - |
| $\omega_m(b,a)$ | value of metric $m$ for reference town $b$ and town $a$ | - | - |
| $\Omega_m(b,T)$ | weighted sum of metric $m$ for reference town $b$ over the set of towns $T$ | - | - |
| $\eta_m(b)$ | $\frac{\Omega_m(b,T')}{\Omega_m(b,T)}$ | - | proportion |
| $\epsilon_m(b)$ | watershed enrichment of metric $m$, that is, $\frac{\eta_m(b)}{c(b)}$ | - | proportion |
| $\Omega_{IBD}(b,a)$ | total IBD between individuals in town $a$ and town $b$ | O | centiMorgans |
| $\Omega_{mig}(b,a)$ | number of migrants from town $a$ to town $b$ (equivalent to $\delta_{a\to b}$) | O | individuals |
| $\Omega_{coal}(b,a)$ | overlap of relative cross coalescence between individuals in town $a$ and town $b$ (equivalent to $\gamma^{ba}$) | E | probability for time of first coalescence for all pairs of lineages |

#### Supporting tables and figures

| **Cohorts** | | | | |
|---|---|---|---|---|
| Quebec (Cartagene) | Quebec (Genizon) | France (Suvimax) | GBR (1kGP) | Total |
| 12,064 | 9,004 | 2,276 | 91 | 23,435 |
| **Inferred Ancestry** | | | | |
| FC (genealogy) | FC (inferred) | non-FC (inferred) | Europe | Total |
| 4,882 | 15,569 | 617 | 2,367 | 23,435 |

Table S1: Sample sizes of genotyped cohorts and their ancestries. A total of 20,451 individuals genealogically linked or genetically inferred French-Canadian (FC) ancestry are used for visualizing population structure using PCA and UMAP.

| **South-East** |
|---|
| 1, 4, 5, 6, 7, 13, 26, 38, 42, 69, 73, 74, 83, 84 |
| **South-West** |
| 9, 11, 12, 24, 30, 31, 32, 33, 34, 40, 46, 47, 48, 64, 65, 66, 81, 82 |
| **West** |
| 16, 17, 22, 29, 35, 44, 49, 53, 56, 72, 79, 85, 86 |
| **North-West** |
| 2, 14, 27, 50, 59, 60, 61, 62, 76, 80 |
| **Central** |
| 3, 15, 18, 19, 23, 28, 36, 37, 41, 43, 45, 63, 87 |
| **Isle-of-France** |
| 75, 77, 78, 91, 92, 93, 94, 95 |
| **East** |
| 8, 10, 21, 25, 39, 51, 52, 54, 55, 57, 58, 67, 68, 70, 71, 88, 89, 90 |

Table S2: French administrative departments within each of the seven regions.

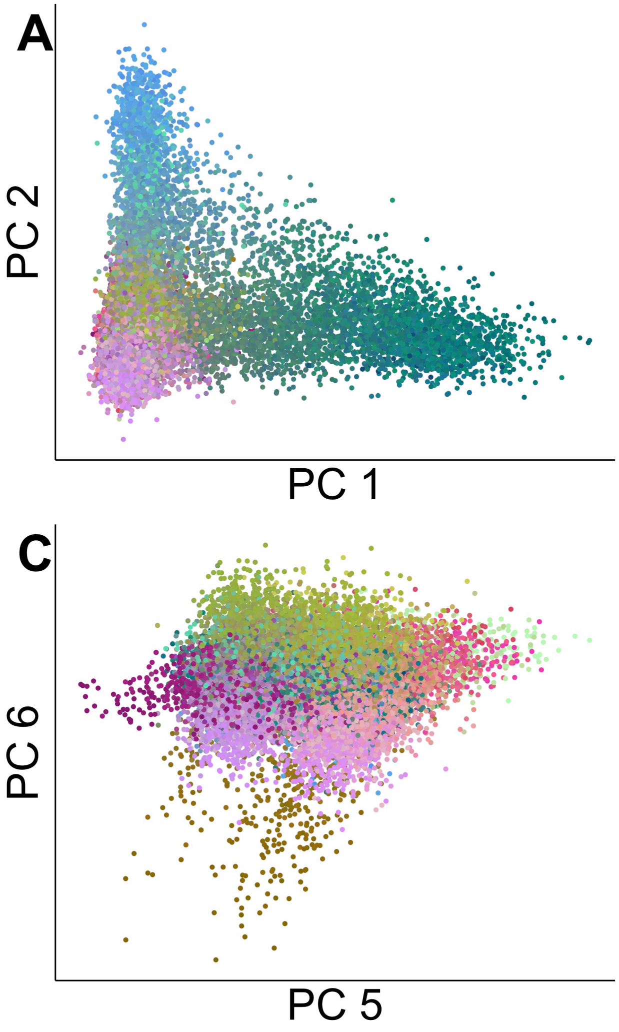

Supplementary Figure S1: **Principal Component Analysis of French Canadians** (**A-D**) The top eight principal components of the genotype data included in the analyses.

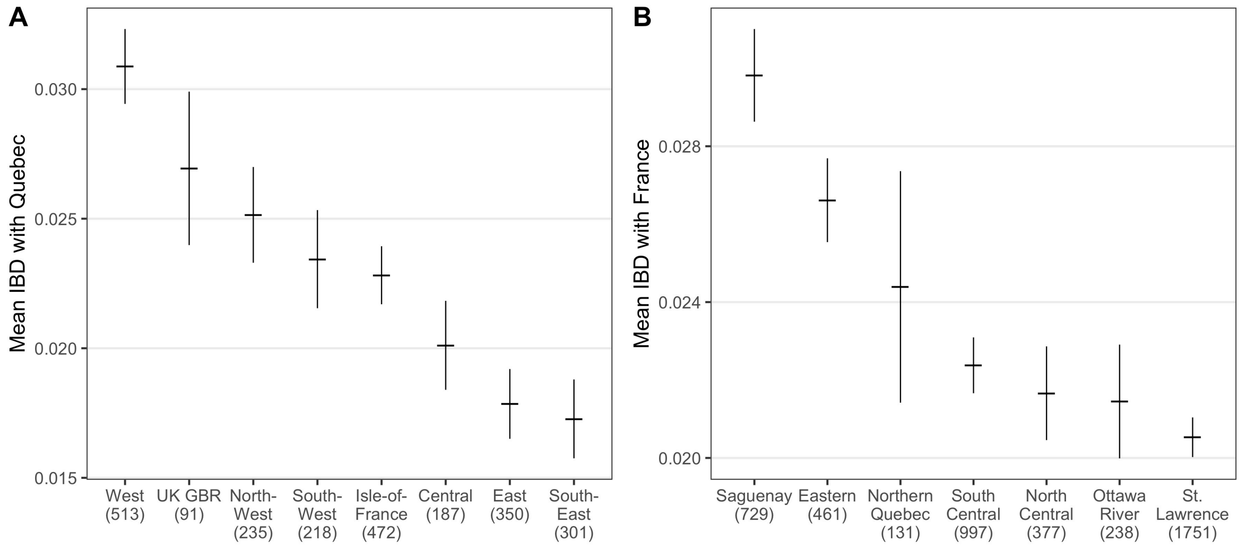

Supplementary Figure S2: **Rates of identity by descent in different regions.** The average IBD sharing between individuals living in Quebec, France, and England. The error bars in these plots indicate the standard error of the mean length (cM) of IBD sharing across individuals from the region indicated on the x-axis. As expected, regions with low sample sizes (indicated on the x-axis) have larger standard errors. (**A**) Individuals from Quebec have higher rates of IBD with individuals from Western and Northern France than with individuals from Central and Southern France. We note that in addition to having a small sample size and large error bars, the GBR cohort is less comparable to French cohorts because their genomic data are generated using different platforms. (**B**) Individuals from France have higher rates of IBD with individuals from Saguenay and Eastern Quebec than with individuals from the Ottawa River and Central Quebec.

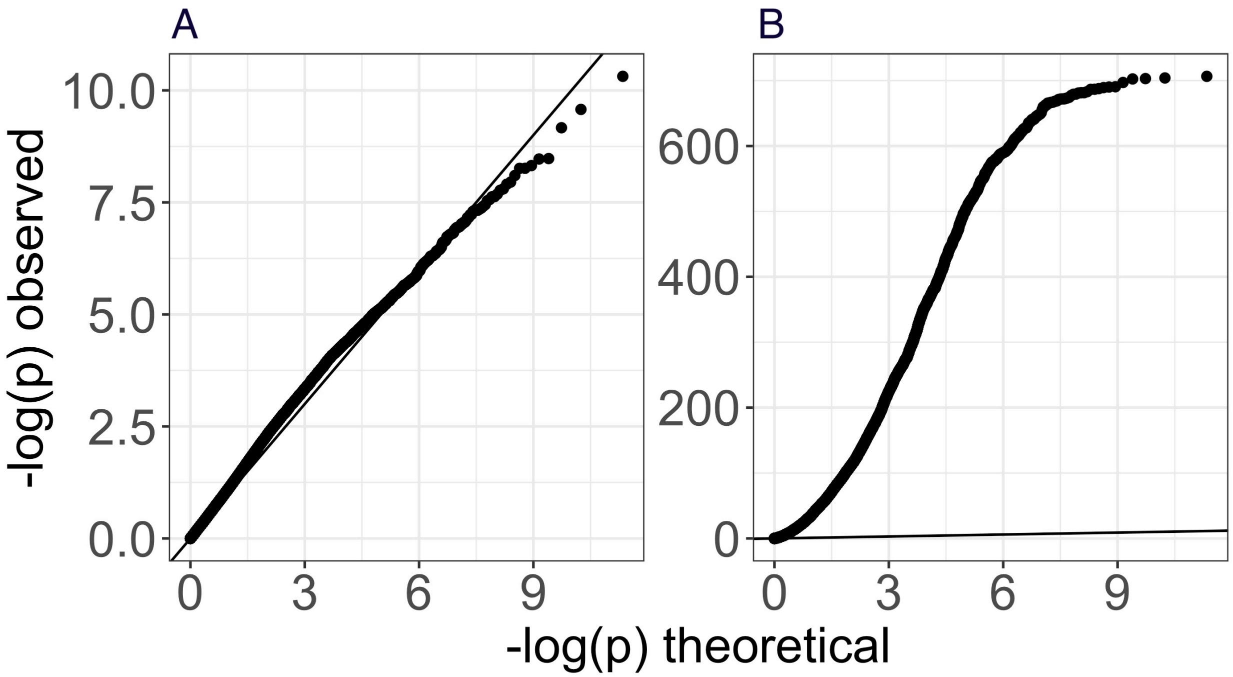

Supplementary Figure S3: **Comparison of Quebec, French and British regions using $F_4$-statistics (A)** All 441 combinations of regions in Quebec and Europe (France and Britain) were compared using an $F_4$-statistic: $F_4$(Qc1, Qc2, Eu1, Eu2). The QQ plot shows no enrichment of significant $p$-values, consistent with the null hypothesis that European population structure is not strongly preserved in Quebec. **(B)** We computed complementary $F_4$ statistics comparing regions in Europe and regions in Quebec, to other regions in Europe and Quebec ($F_4$(Qc1, Eu1, Qc2, Eu2)), and find that all are significantly different, confirming that there are enough genetic differences between these populations to identify $F_4$ statistic signal.

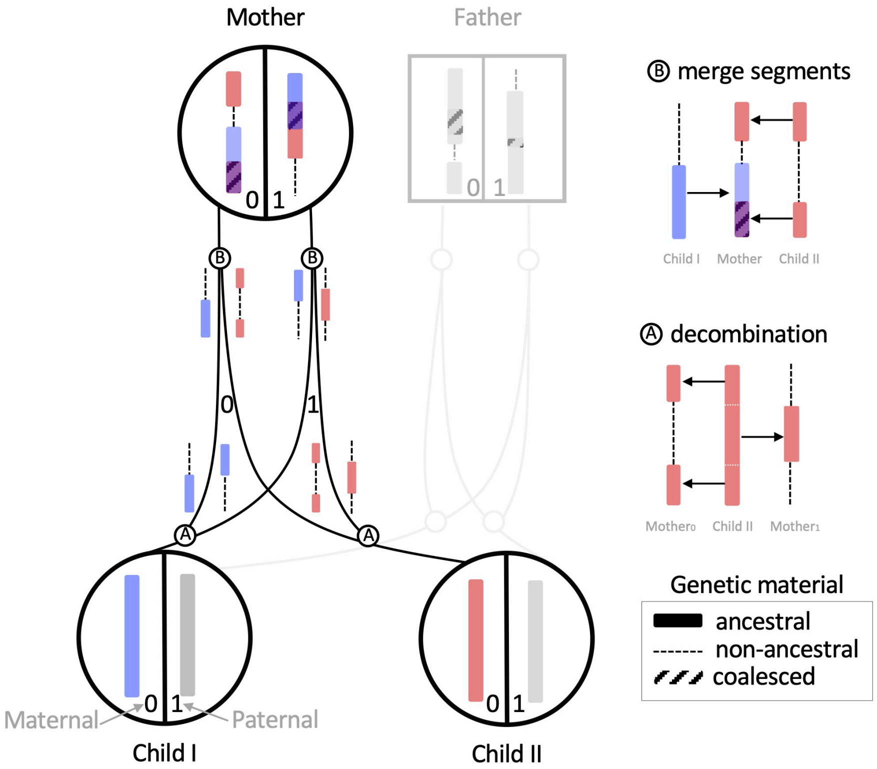

Supplementary Figure S4: **Illustrating decombination** (**A**) Ancestry simulations in backwards time *decombine* genetic material from children based on a user specified recombination map. (**B**) Decombined segments are then merged in a common ancestor. Not all of the genetic material contributed to children coalesces with other ancestral material.

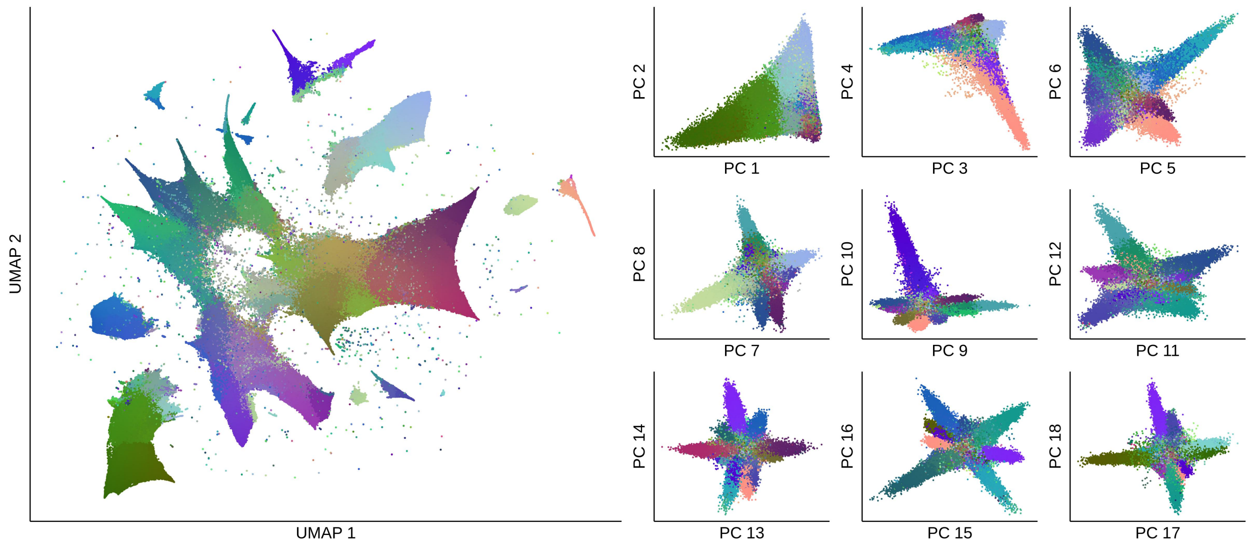

Supplementary Figure S5: **Visualizing 1.4 Million simulated French Canadian genomes (left)** A UMAP of the simulated genotype data. (**right**) The top eighteen principal components of the simulated genotype data. Colours were generated from a three dimensional UMAP through converting each $x, y, z$ coordinate into an RGB value unique to each individual (see Supplementary Methods 3.2).

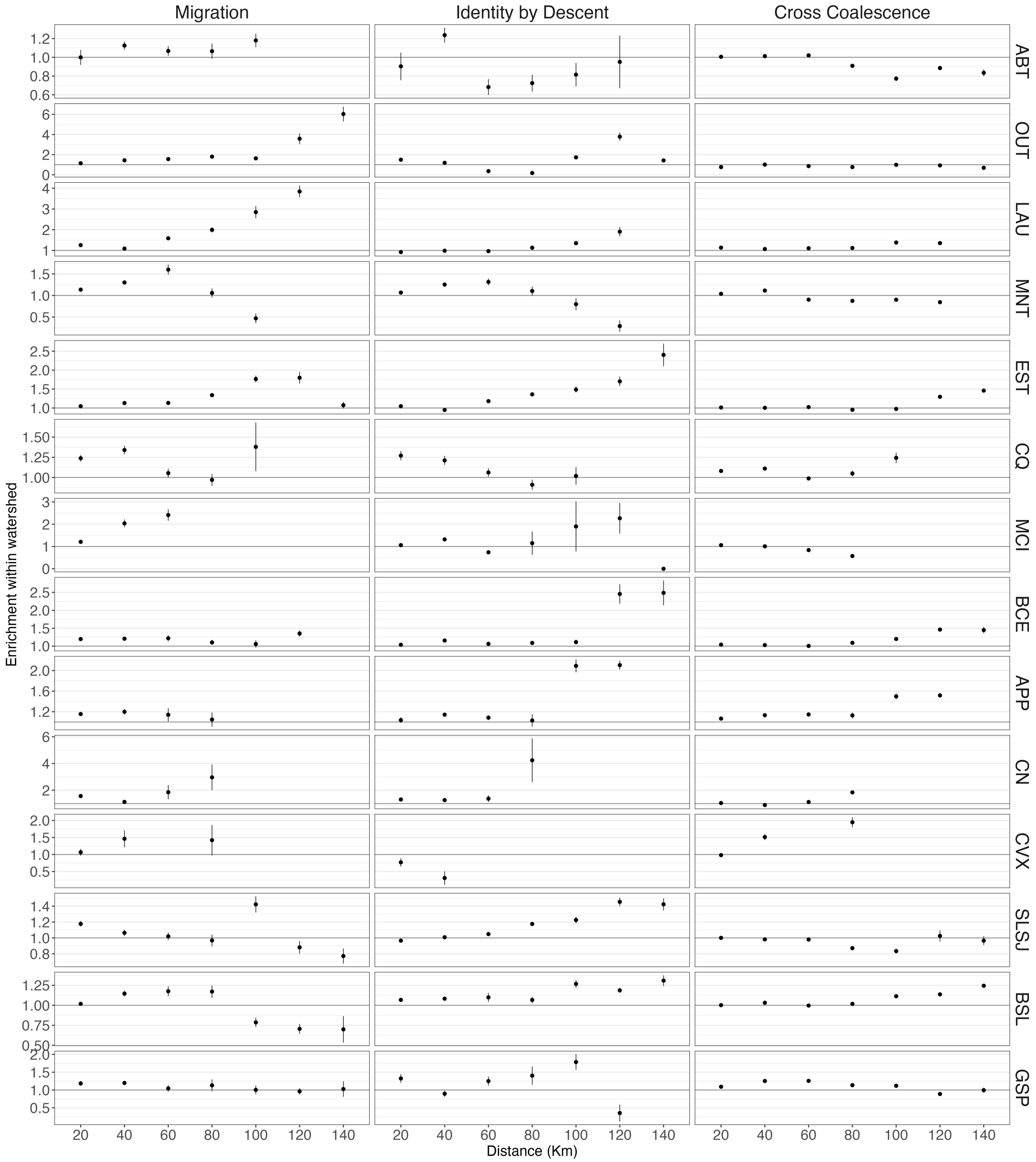

Supplementary Figure S6: **Historical migrations that define regional population structure** Watersheds influence migrations, genetic and genealogical relatedness. Rows of panels indicate a separate region's watershed enrichment comparing the migration rates IBD rates and cross coalescence rates between pairs of individuals in towns within and without the same watershed. Note the y scale along rows are consistent, but across rows are scaled separately.

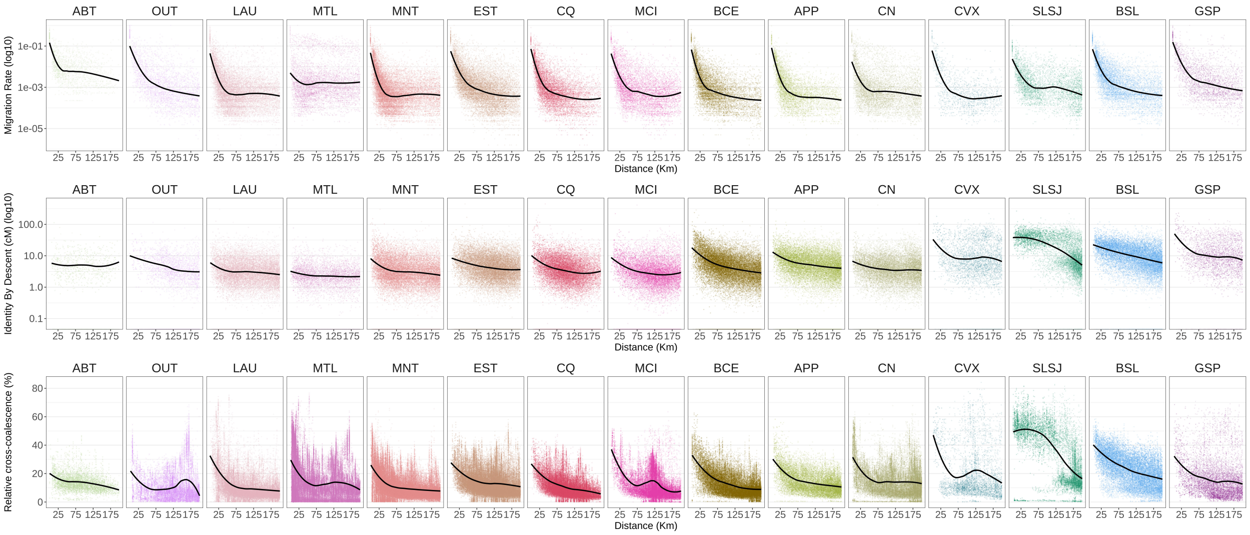

Supplementary Figure S7: **Migration rates, Identity by descent rates, and relative cross coalescence rates decay with distance for each region in Quebec.** The y axis is $\omega_m(b, a)$ for all metrics $m$ for all reference towns $b$ and distal towns $a$. The x axis is the distance in kilometres between reference towns $b$ and distal towns $a$. Rows of panels show the decay of a single metric across regions in Quebec. Columns correspond to the regions defined in S16. The solid black line indicates a loess fit line using local polynomial regression fitting for each panel separately.

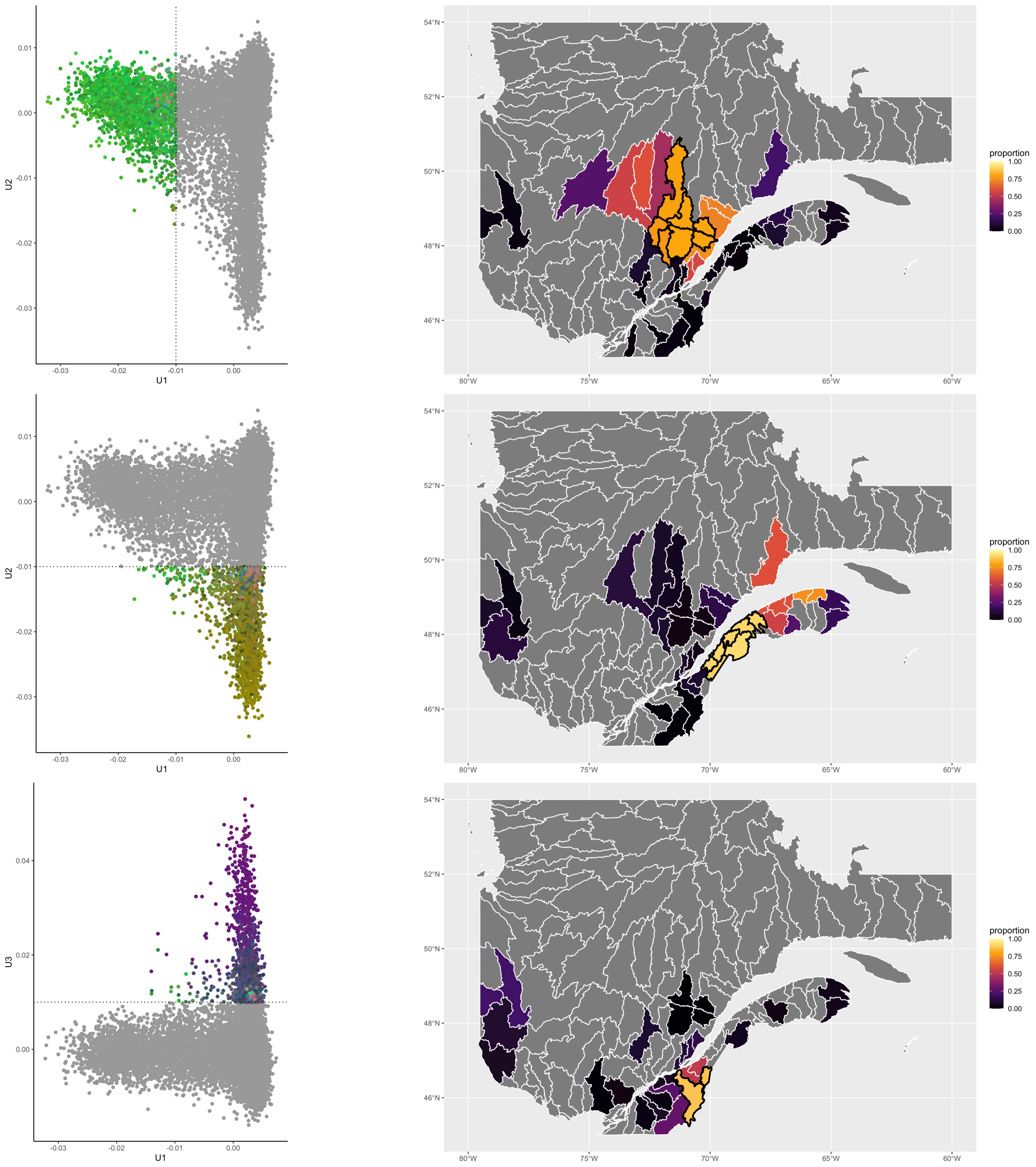

Supplementary Figure S8: **Selecting watersheds with individuals driving principal components** We define a threshold of 0.01 along each of the top three principal components and compute the fraction of individuals in each watershed that are beyond this limit. We choose watersheds with more than 75% of individuals beyond the threshold (black contours).

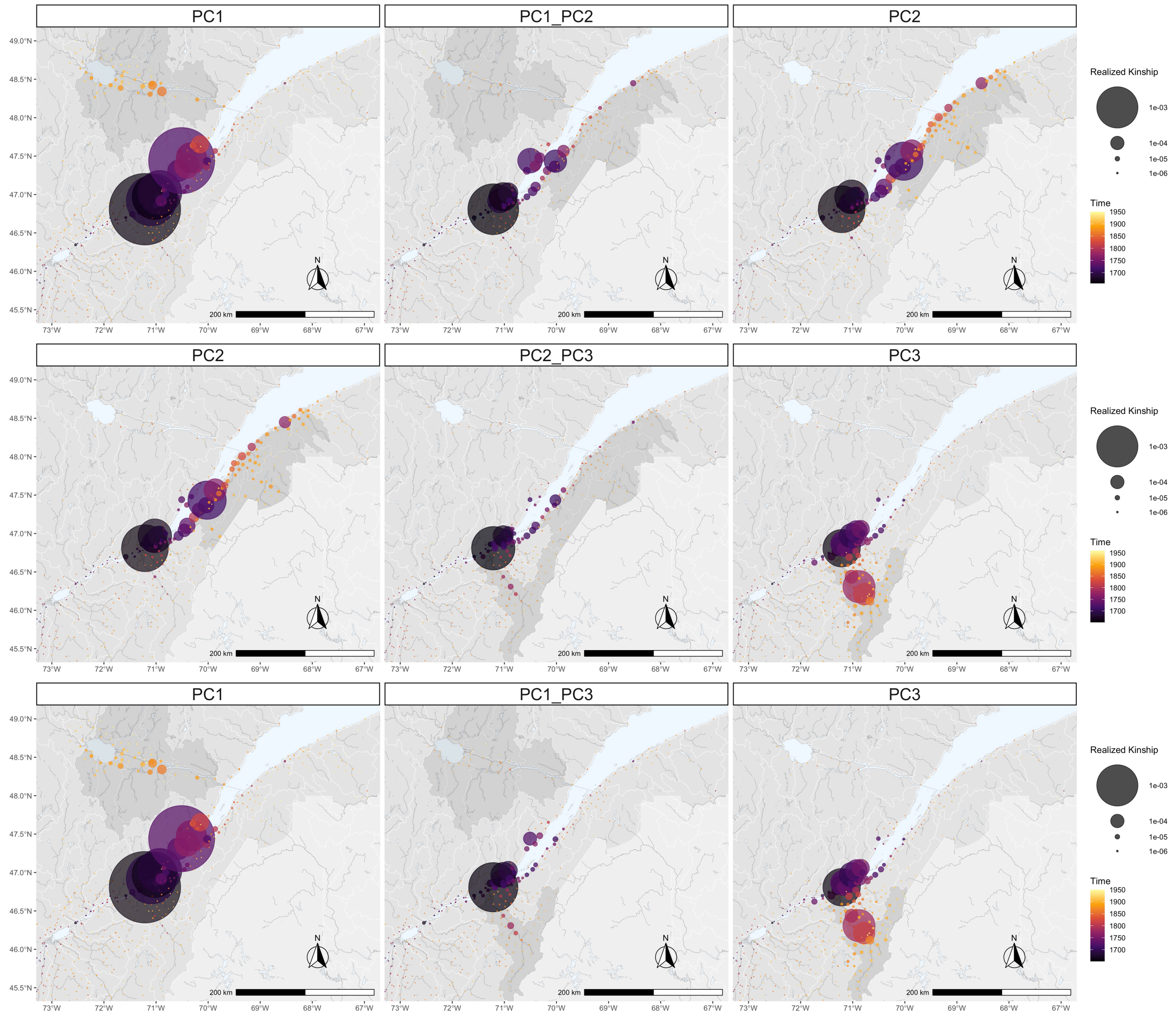

Supplementary Figure S9: **Cross coalescence rates** For each pair of regions defined in Figure S8, we compute the cross coalescence rates by ascending the genealogies of individuals in both regions. The central panels represent the between region cross coalescence rate, whereas the first and last column of panels represent the within region coalescence rate. We find that in all three cross coalescence rates, there is a common root in the region around Quebec City.

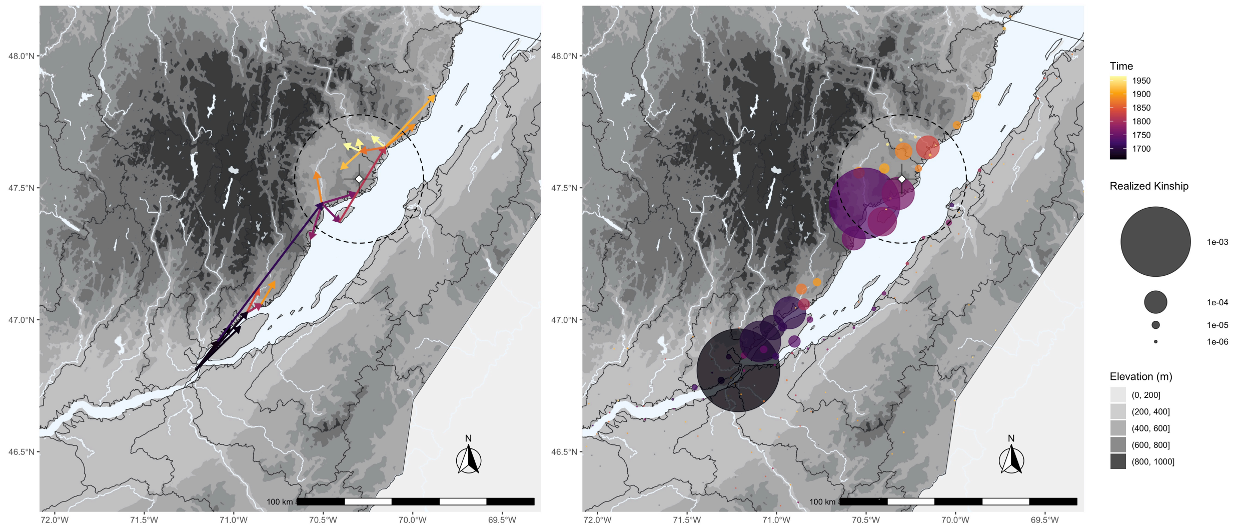

Supplementary Figure S10: **Charlevoix astrobleme impact on population structure** For present day individuals living in Charlevoix (highlighted area), we show **(A)** the major migratory axes as well as **(A)** the location, timing and stringency of population bottlenecks measured by realized kinship. The epicentre of the astrobleme is marked with a cross and the radius of the ancient meteor crater is indicated with a dotted line.

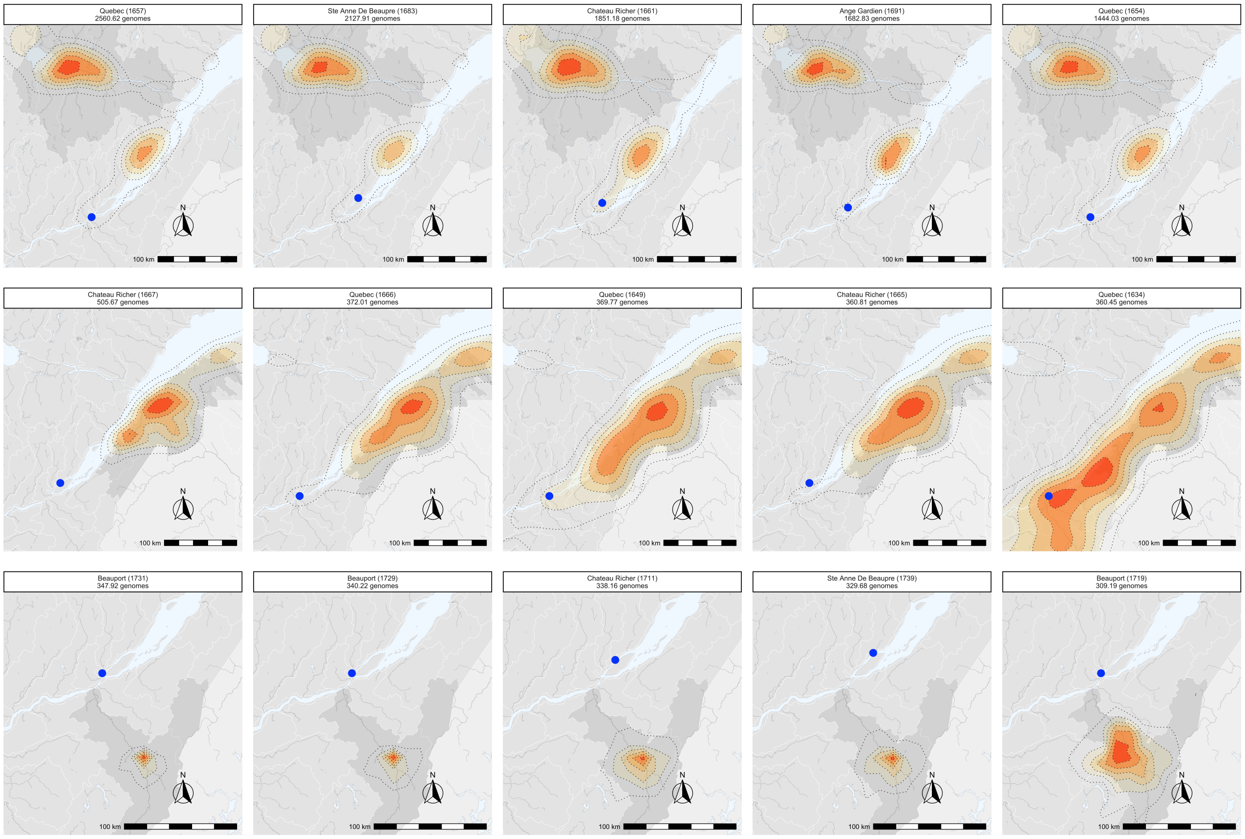

Supplementary Figure S11: **Dispersal range of the top five contributors to each region** For each region **(rows)** enriched in individuals driving each of the top three principal components, we show the dispersal range of the top five super-founders **(columns)**. Even though there is some variation between the dispersal ranges of the super-founders of a region, they broadly cover the same geographic regions. The scaling of the heat maps are not comparable between plots as they were computed separately.

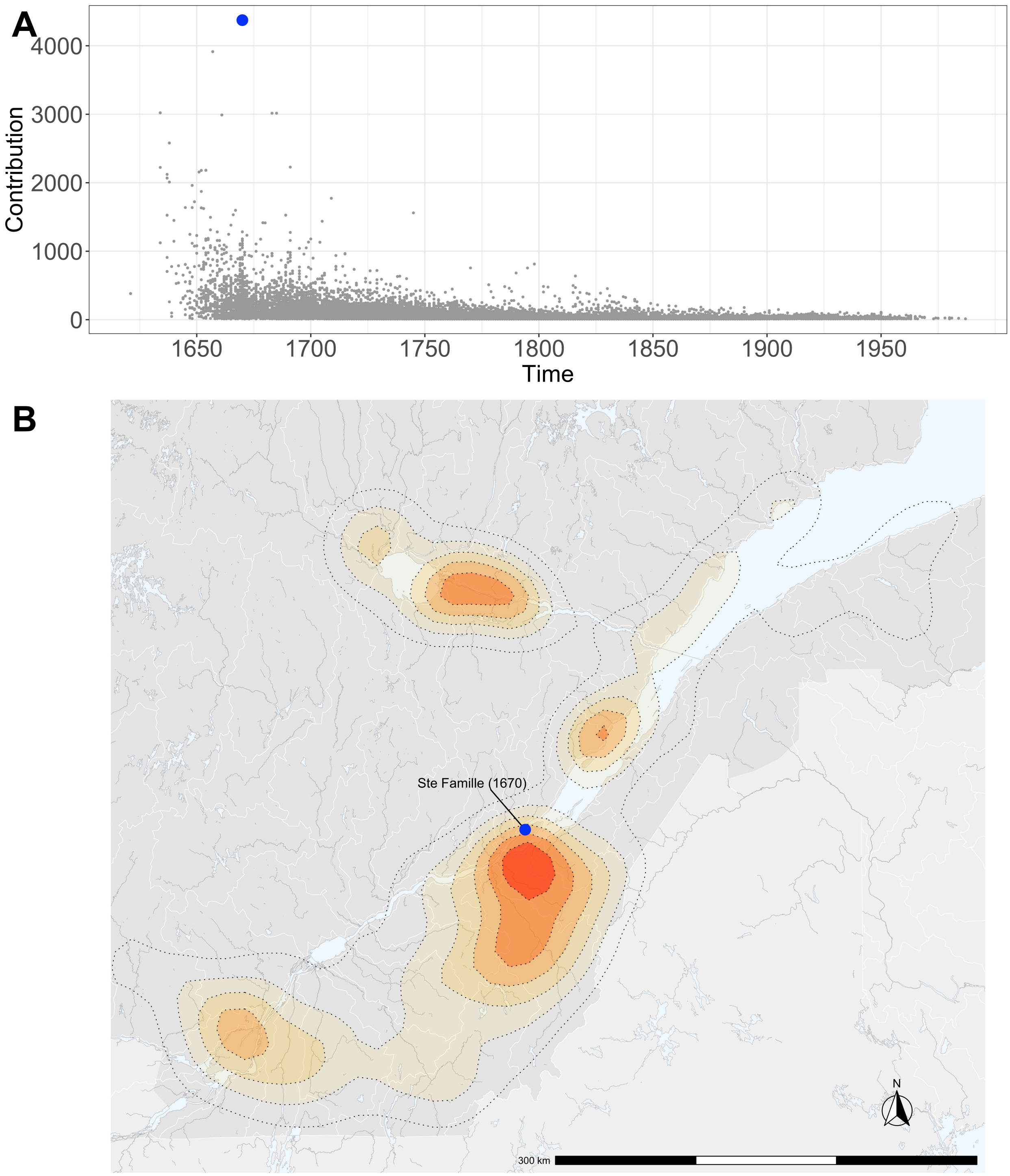

Supplementary Figure S12: **Range dispersal for the top contributing ancestor A** The total genetic contribution of all ancestors in the pedigree. A blue dot highlights the ancestor with the greatest contribution to individuals in the pedigree. **B** The range dispersal of the historical individual with the greatest contribution to probands.

| Parameter | Specification | Citation |
|---|---|---|
| demographic model | European ancestry in two population out-of-Africa model | (*31*) |
| coalescent model | Hudson | (*59*) |
| mutation rate beyond known pedigree | $3.6210^{-8}$ | (*51*) |
| recombination map | GRCh37 hapmapII genetic map | (*50*) |

Table S3: The model parameters used to simulate the ancestry of individuals in the French-Canadian pedigree.

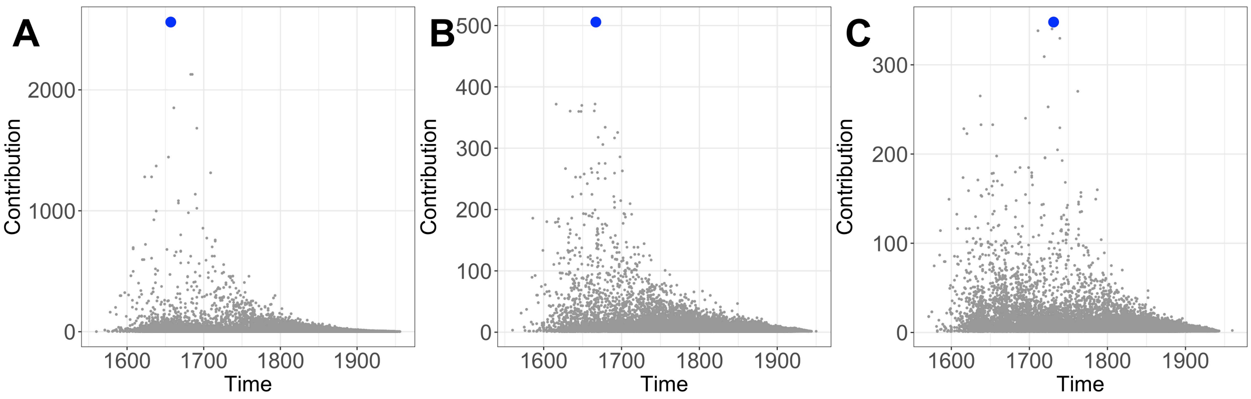

Supplementary Figure S13: **Genetic contribution of ancestors** The genetic contribution of each ancestor to probands living in each of the three regions enriched in the top three principal components. The x axis places each ancestor in time based on their marriage date. The ancestor contributing the most genetic material to each region is indicated by a blue dot.

| PC1 | PC2 | PC3 |
|---|---|---|
| 2561 | 122 | 87 |
| 2128 | 93 | 60 |
| 1851 | 83 | 64 |
| 1683 | 91 | 15 |
| 1444 | 74 | 16 |
| 1314 | 79 | 14 |
| 1137 | 51 | 48 |
| 984 | 50 | 21 |
| 124 | 506 | 12 |
| 134 | 372 | 16 |
| 146 | 370 | 37 |
| 119 | 361 | 15 |
| 201 | 360 | 171 |
| 119 | 360 | 19 |
| 118 | 334 | 23 |
| 66 | 326 | 6 |
| 94 | 317 | 21 |
| 8 | 4 | 348 |
| 8 | 4 | 340 |
| 9 | 4 | 338 |
| 9 | 4 | 330 |
| 9 | 4 | 309 |
| 6 | 3 | 270 |
| 520 | 87 | 265 |
| 7 | 3 | 253 |

Supplementary Figure S14: **Contributions of regional super-founders** For each region enriched in individuals driving each of the top three principal components, we identify the top ten 'super-founders' based on their realized kinship to each region. We also compute the contributions of these ancestors to individuals living in the other two regions to see how much overlap there between their descendants. We find that each of the super-founders disproportionately contributes to a single region.

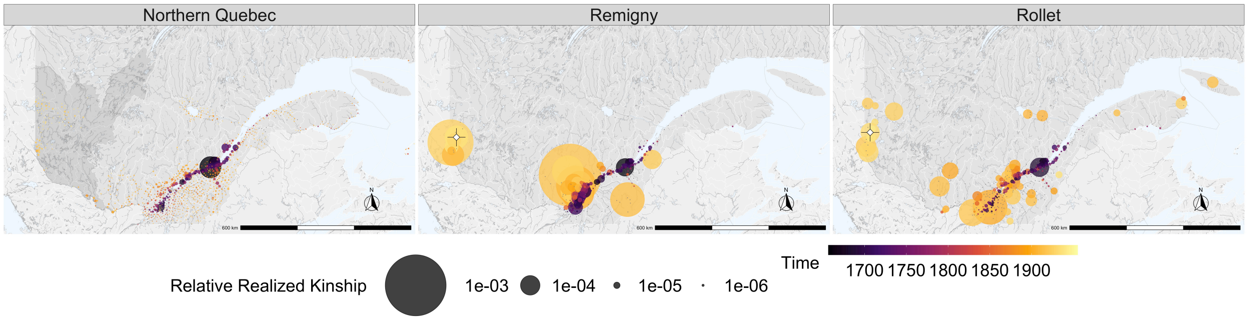

Supplementary Figure S15: **Realized kinship for Remigny and Rollet** Northern Quebec (left panel) shows no major regional founding events other than in Quebec City in the 16th century. When we consider each town separately, bottlenecks become more apparent. The towns of Remigny (centre panel) and Rollet (right panel) have recent founding events from different regions in Quebec despite being twenty kilometres away from each other.

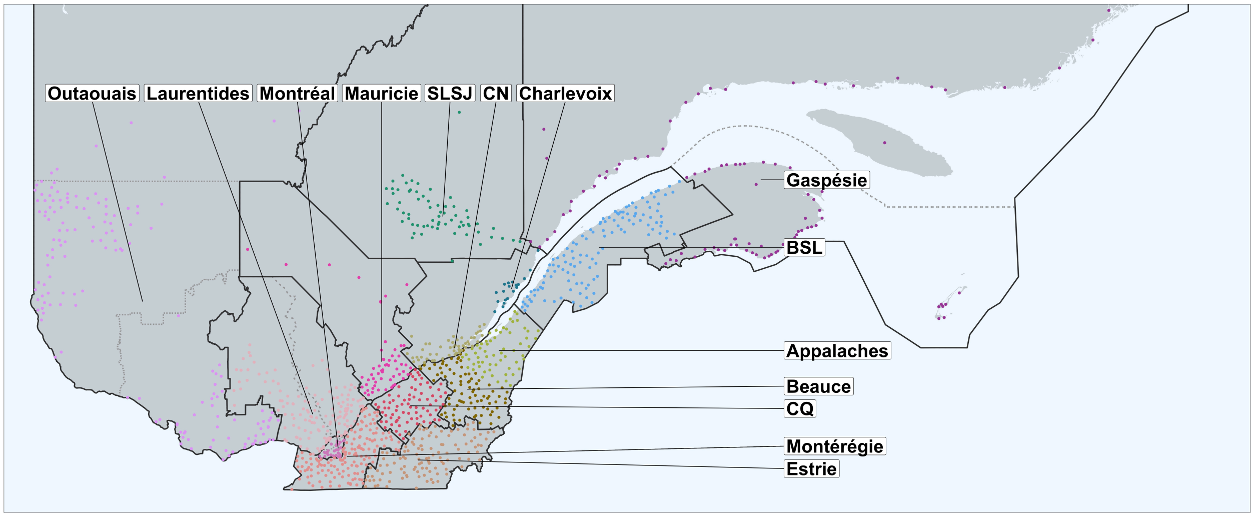

Supplementary Figure S16: **Administrative boundaries used to generate genealogy flow plot** The genealogy flow plot in Figure 1 was generated based on Quebec administrative boundaries. We modified the groupings of some regions because they had similar demographic history or small population sizes. The regions of Nord-du-Québec, Abitibi-Témiscamingue, and Outaouais were clumped into a single group. The regions of Laval and Montréal were clumped together. The regions of Lanaudière and Laurentides were clumped together. The regions of Cote-Nord, Gaspésie and Iles-de-la-Madeleine were also clumped together. In addition, we separated two administrative regions with demographic histories that we sought to visualize separately. The region of Charlevoix was separated from Capitale-Nationale and the region of Chaudière-Appalaches was separated into Beauce and Appalaches.

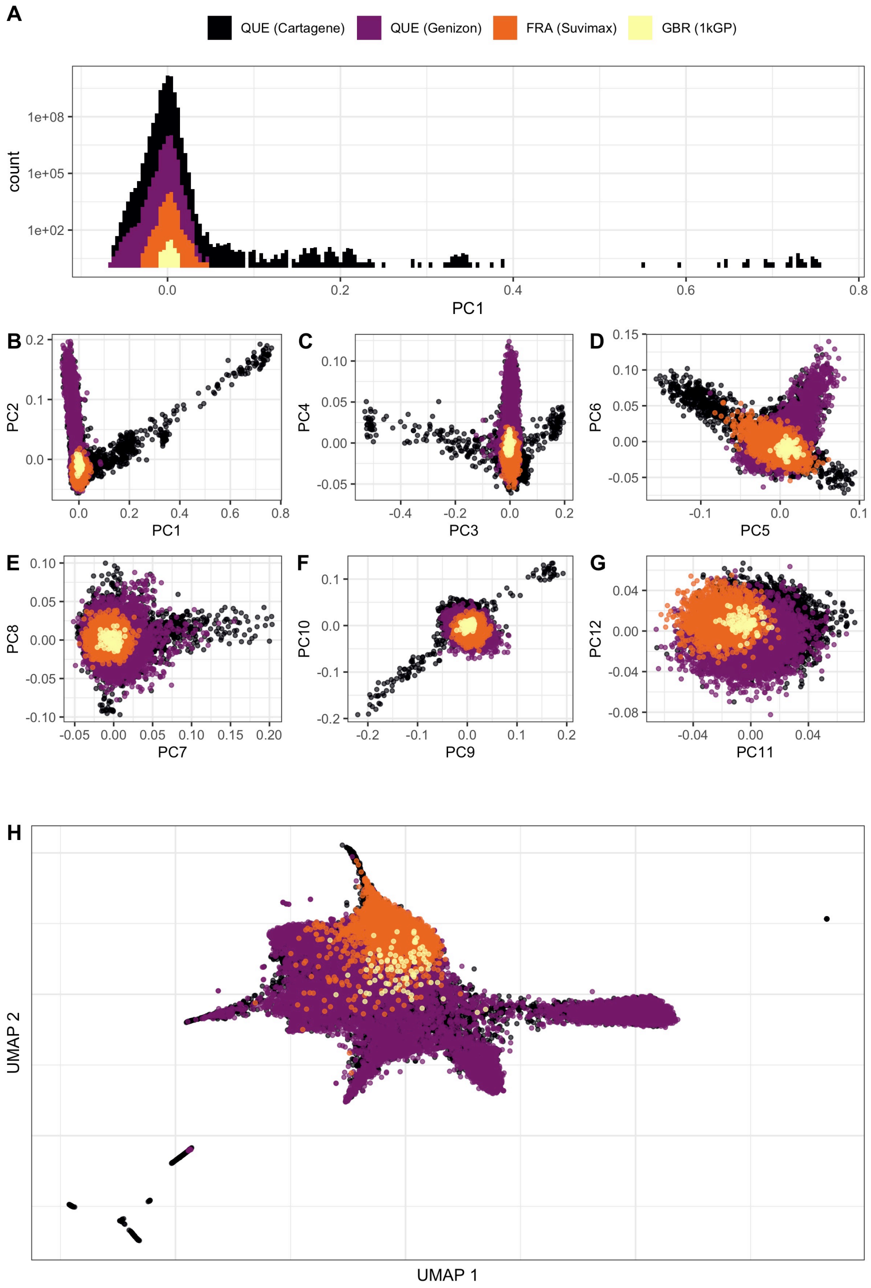

Supplementary Figure S17: **Visualization of the PCA and UMAP analyses of the complete genotype dataset.** **A** Is the distribution of samples along the first principal component using all samples form all cohorts. We colour each cohort separately to highlight any possible batch effects or population differentiation. **B-G** Are the top twelve principal components of all samples form all cohorts. **H** Is a UMAP of these samples using the top ten principal components.

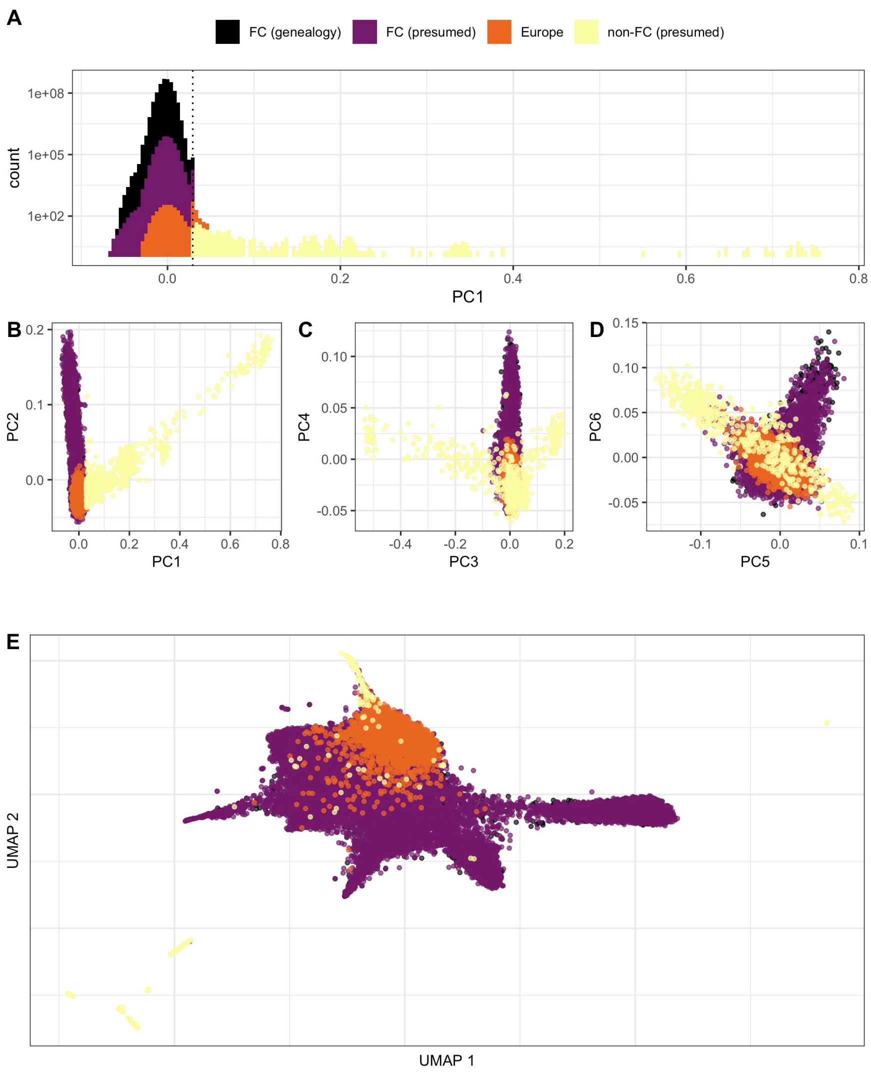

Supplementary Figure S18: **Visualization of the thresholds used to define presumed French Canadian ancestry. A** Is the distribution of samples along the first principal component using all samples form all cohorts. We colour samples based on their ancestries. In this case, we consider samples from France and from Britain as European, and samples from Quebec as either genealogically confirmed FC ancestry, presumed FC ancestry using a threshold based on the maximum value along the first principal component of genealogically linked individuals, and non-FC ancestry for individuals beyond this threshold. **B-G** Are the top twelve principal components of all samples form all cohorts coloured based on presumed ancestry. **H** Is a UMAP of these samples using the top ten principal components coloured based on presumed ancestry.
